# Repeat elements organize 3D genome structure and mediate transcription in the filamentous fungus *Epichloë festucae*

**DOI:** 10.1101/339010

**Authors:** David J Winter, Austen RD Ganley, Carolyn A Young, Ivan Liachko, Christopher L Schardl, Pierre-Yves Dupont, Daniel Berry, Arvina Ram, D Barry Scott, Murray P Cox

## Abstract

Structural features of genomes, including the three-dimensional arrangement of DNA in the nucleus, are increasingly seen as key contributors to the regulation of gene expression. However, studies on how genome structure and nuclear organization influence transcription have so far been limited to a handful of model species. This narrow focus limits our ability to draw general conclusions about the ways in which three-dimensional structures are encoded, and to integrate information from three-dimensional data to address a broader gamut of biological questions. Here, we generate a complete and gapless genome sequence for the filamentous fungus, *Epichloë festucae*. Coupling it with RNAseq and HiC data, we investigate how the structure of the genome contributes to the suite of transcriptional changes that an *Epichloë* species needs to maintain symbiotic relationships with its grass host. Our results reveal a unique “patchwork” genome, in which repeat-rich blocks of DNA with discrete boundaries are interspersed by gene-rich sequences. In contrast to other species, the three-dimensional structure of the genome is anchored by these repeat blocks, which act to isolate transcription in neighbouring gene-rich regions. Genes that are differentially expressed in planta are enriched near the boundaries of these repeat-rich blocks, suggesting that their three-dimensional orientation partly encodes and regulates the symbiotic relationship formed by this organism.

## Introduction

The advent of high throughput sequencing technologies has revolutionized eukaryotic genomics. Thousands of draft genome sequences have been produced, facilitating their use by an ever-larger group of researchers. However, most of these genome sequences are incomplete and highly fragmentary. A major cause of this incompleteness is that repetitive elements, which are common in many eukaryote genomes, are difficult to assemble with short sequencing reads and short-read genome assemblies therefore frequently break at repeat regions (Schatz et al. 2010). Although fragmentary genome sequences may contain most of the protein coding genes, the lack of high-quality genome sequences for most taxa severely limits our ability to examine the effect of larger-scale genome structure on the regulation of those genes, and the contribution of these higher-level structures to genome evolution.

Species of the genus *Epichloë* are ascomycete fungi that form intimate symbiotic relationships with cool season grasses. In order to maintain these relationships, these fungi must express a suite of genes (Eaton et al. 2010, 2011, 2015) that encode proteins that mediate interactions with their host and catalyse the production of bioprotective alkaloid compounds (Tanaka et al. 2012; Scott et al. 2010). *Epichloë* species can have profound effects on their hosts, including resistance to herbivory by mammals (Schardl et al. 2007) and insects (Rowan & Gaynor 1986; Clay & Cheplick 1989), resistance to nematodes (Kimmons 1990) and *non-Epichloë* competitor fungi (Christensen 1996), and increased drought tolerance (Malinowski & Belesky 2000; Bayat et al. 2009; Arachevaleta et al. 1989). The bioprotective properties conferred by *Epichloë* species provide substantial economic benefits (Kauppinen et al. 2016; Saikkonen et al. 2016; Lugtenberg et al. 2016), as many agriculturally important grasses form relationships with *Epichloë* species.

The intimate symbiotic relationship and the powerful techniques available to interrogate the *Epichloë-grass* interaction make this the most well-developed system in which to study an above-ground interaction between a fungus and a plant (Tanaka et al. 2012; Schardl 2001). Because *Epichloë* species can be genetically manipulated, maintained in culture, and used to create experimental infections, the interaction can be studied with the full range of modern molecular techniques (Young et al. 2005; Clarke et al. 2017; Eaton et al. 2015). Deep RNA sequencing of infected grass tissues makes it possible to measure the expression of *Epichloë* genes *in planta*. Comparing the expression of *Epichloë* genes between culture and *in planta* conditions and between wild type and symbiosis-deficient strains has led to the identification of a suite of genes required to establish and maintain the symbiosis (Eaton et al. 2010, 2011). Despite considerable efforts, the precise means by which the expression of these genes is regulated remains elusive.

High throughput chromosome conformation capture (HiC) and related technologies have made it possible to study the three dimensional organization of genomes and examine chromatin states at a genome-wide scale (Lieberman-Aiden et al. 2009; Bonev & Cavalli 2016). Studies taking advantage of these technologies have revealed the means by which nuclear organization is encoded and maintained in various lineages, and demonstrated the important roles these structures play in regulating gene expression across the tree of life (Li et al. 2012; Duan et al. 2010; Cubeñas-Potts et al. 2017). At present, however, little is known about nuclear organization in filamentous fungi and our ability to examine the three dimensional structure of *Epichloë* genomes is limited by the fact the only reference sequences available for this genus are highly fragmentary (Schardl et al. 2013). These unfinished genomes make it difficult to analyses HiC data at a genome-wide scale.

Here we combine a range of modern sequencing techniques, including long read sequencing and HiC, to investigate the role that genome structure plays in gene regulation in *Epichloë*. Long read sequencing has allowed us to generate a gapless telomere-to-telomere assembly for *E. festucae* strain Fl1, a model system for *Epichloë* research (Scott et al. 2012). Our sequences reveal a remarkable ‘patchwork’ genome, in which regions with very high repeat-density are interlaced with almost repeat-free sequences. Using chromosome conformation to investigate the three-dimensional structure of the genome, we show that these repeat-rich blocks mediate genome folding in the nucleus. This genome structure contributes to the modulation of gene expression, notably for those genes that are strongly differentially expressed *in planta*.

## Results

### Assembly of a complete and high quality *E. festucae* genome sequence

Existing genome sequences for *E. festucae* are highly fragmented, each containing more than 1000 contigs with N_50_ < 90 kb (Schardl et al. 2013). To generate a high quality contiguous reference assembly, we used PacBio and Illumina shot-gun sequencing combined with high throughput chromosome conformation capture (HiC) data. We obtained high quality, high molecular weight genomic DNA from *E. festucae* strain Fl1 (Moon et al. 1999; Leuchtmann 1994), and subjected this to sequencing. After filtering, PacBio long-read sequencing yielded 948,767 subreads with an N_50_ of 10,019 nucleotides. The approximately 7.2 billion total bases sequenced represent a mean coverage of 211x for the genome. We generated separate assemblies from these reads using the PacBio assemblers Canu and HGAP. The Canu assembly was the most contiguous, with 35 Mb contained in 21 contigs. Four of these contigs were capped by telomeric repeats on both ends, suggesting they represent complete chromosomes.

We used HiC data to generate complete assemblies of the remaining chromosomes. Contigs were joined using HiC reads that map to the ends of different contigs. The HiC scaffolded assembly had a total of seven putative chromosomes, each capped by telomeric repeats on both ends. This assembly contained just 5 gaps, which were manually in-filled using sequence from the HGAP assembly, whose alternative algorithm had assembled across the remaining gaps from the Canu assembly. HGAP contigs that aligned to >10 kb of sequence on the ends of two Canu contigs were used to fill gaps (Figure 1a).

**Figure 1.**
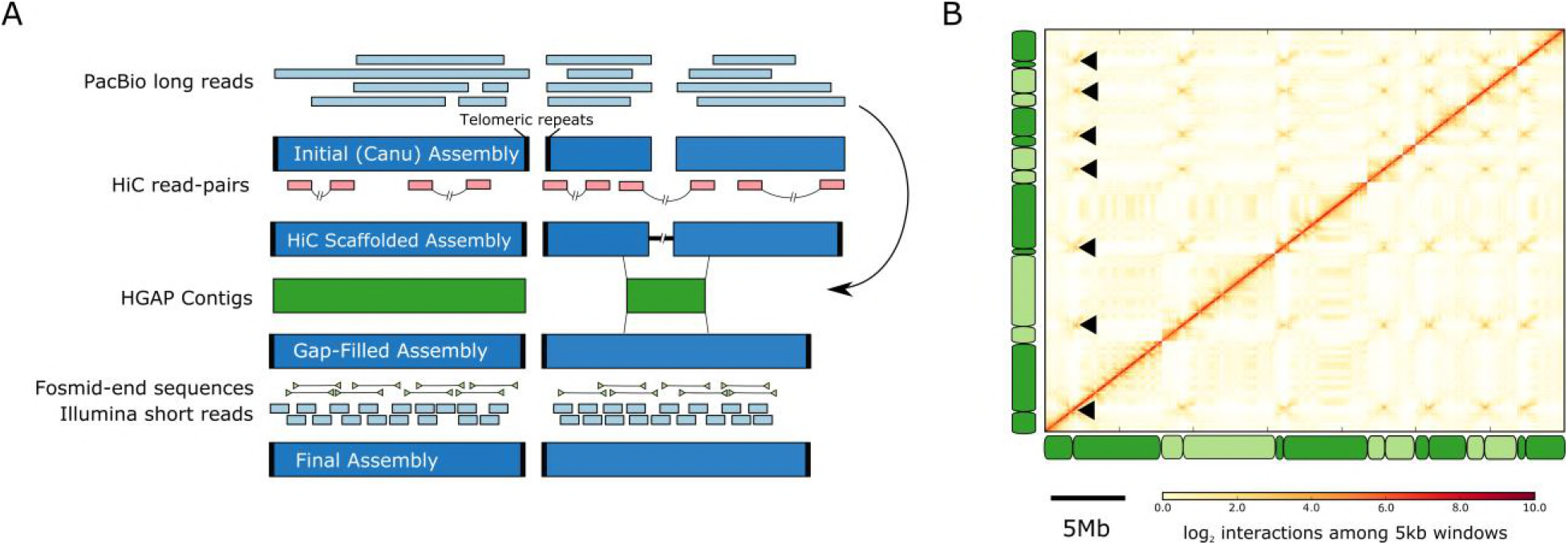
An Iterative process generated a complete and gapless genome assembly. **A**: PacBio reads were assembled with Canu. This initial assembly included complete chromosomal sequences (capped by telomeric repeats) and smaller sub-chromosomal contigs. HiC data provided information about the physical proximity of genomic regions, allowing the joining of contigs into scaffolds. Gaps in this scaffolded assembly were filled using an independent genome assembly made with the HGAP assembler, which uses a different algorithm, from the same PacBio reads as in the initial assembly. Finally, Illumina short reads and Sanger end sequences of fosmids were used to correct base-errors and test the structural integrity of the final assembly. **B** Genome-wide HiC contact map derived from cells in culture. Each element of the matrix reflects the frequency of contacts between two 5 kb windows in the genome assembly. The positions of chromosomes in the whole genome data are indicated along the left and bottom edges, with adjacent chromosomes indicated by different shading. Black arrowheads mark the putative centromeres, which are indicated by characteristic cross-like patterns.

Two contigs remained that had a telomere at one end and part of the ribosomal RNA gene repeats (rDNA) at the other. In fungi the rDNA is organized into tandem repeats with 10s-100s of copies, each with near-identical sequences around 10 kb in length (Ganley & Kobayashi, 2007), precluding its complete assembly even with PacBio sequences. We also found several small, high-coverage contigs containing just rDNA sequence. We generated a consensus rDNA sequence from these, and joined the two large contigs with three full rDNA units, starting by convention with 18S, with contig ends trimmed to retain partial flanking units observed at the rDNA locus boundaries. The ratio of average rDNA coverage versus average genome-wide coverage of Illumina paired-end sequencing data, suggests that the rDNA is present in approximately 21 copies in Fl1.

A contact map derived from the HiC data supports the integrity of our genome assembly (Figure 1B). The strongest signal in the map is the diagonal, as expected if the majority of contacts are occurring with linearly-adjacent genomic regions. We also used the HiC data to estimate the positions of centromeres, which appear as regions of high inter-chromosomal interaction from centromere-to-centromere contacts (Funabiki et al. 1993; Mizuguchi et al. 2014). The existence of six non-self signals for each of our assembled chromosomes is further support for Fl1 containing seven nuclear chromosomes, and establishes the putative centromere position for each chromosome to within 10 kb. No single repeat family occurs at every putative centromere, and we were not able to identify any sequence motif associated with these centromeric positions. *Epichloë festucae* may thus be one of several fungal species that do not have single centromere-defining sequence but instead use epigenetic markers (Sanyal et al. 2004).

A total of 6,709 base-level errors, mostly single base indels (96.9%), were polished using the Pilon finisher with 87x coverage Illumina paired-end sequencing as input. Finally, telomeres were trimmed to the nearest canonical sequence and, where necessary, chromosome sequences were reverse-complemented to place the short arm of the chromosome first by convention. The final assembly contains telomere-to-telomere sequence for seven chromosomes and a complete mitochondrial genome. In total, the assembly contains 35,023,692 bases with no Ns or gaps. We tested the quality of this final assembly by mapping Sanger end-sequences of fosmids produced from *E. festucae* Fl1 in a previous project (Schardl et al. 2013) to our reference genome. Of 6,472 read pairs, 99.9% mapped in the expected orientation with a median inferred insert size of 35.3 kb (sd = 4.4 kb), as expected for fosmid data. The only region with aberrant size mapping was the rDNA locus, a result that is expected due to the artificially reduced copy number for this locus in our assembly.

We used the complete reference genome to discover genes that had not been revealed by existing fragmentary genome sequences. By mapping RNAseq reads derived from Fl1 in culture and *in planta* to our complete reference genome, we identified a total of 8,959 transcript-encoding loci. Of these, 8,150 (91%) overlap with the coding sequence from one of the “M3” gene models produced from the closely related *E. festucae* strain E2368 (Schardl et al. 2013). Among the 809 genes that do not correspond to an M3 gene model, 788 (97%) have sequence homology to regions of the fragmented reference genome previously available for *E. festucae* Fl1, and 672 (83%) share homology with a region of the existing fragmentary assembly of strain E2368. This result suggests existing transcriptome assemblies already contain sequences for the majority of protein coding genes. For this reason, we focused our subsequent analyses on how the structural features of the genome revealed by our sequence contribute to the regulation of gene expression.

### The genome is a patchwork of sequences with distinct nucleotide content

The genomes of some filamentous fungi contain distinct regions with unusually low GC content (Testa et al. 2016). The *E. festucae* Fl1 genome is a striking example of this phenomenon. The genome contains alternating blocks of AT-rich sequences and sequences with approximately equal nucleotide composition (Figure 2, Table 1). These blocks are distinct not only in nucleotide composition, but also in their content. The AT-rich regions largely comprise repetitive DNA, with 85% of bases in these regions showing similarity to a known repeat, while only 0.09% of these regions are part of a known gene. In contrast, only 0.8% of bases outside of the AT-rich regions are part of a known repeat, whereas 56% of these bases are part of an exon (Table 1). Hereafter we use the term ‘gene-rich’ to describe the regions of the genome with approximately equal nucleotide content. Consistent with previous reports, we found a diverse range of repetitive elements including LTR retrotransposons, Miniature Inverted Repeat Transposable Elements (MITEs) and DNA transposons (Fleetwood et al. 2011; Schardl et al. 2013). Although most of the repetitive DNA in the *E. festucae* Fl1 genome falls into the AT-rich component of the genome, smaller elements such as MITEs are unique in being predominately found in the gene-rich component (Supplementary Figure S3). Most of AT-rich blocks are comprised of multiple repeats. Repeats within these blocks form complex nested structures, with contiguous repeat copies being interrupted by one or more other repeat sequence (Figure 3A). This nesting of repeats is so widespread that most copies of most repeat families are interrupted by other repeat copies from multiple families (Supplementary Figure S2).

**Figure 2:**
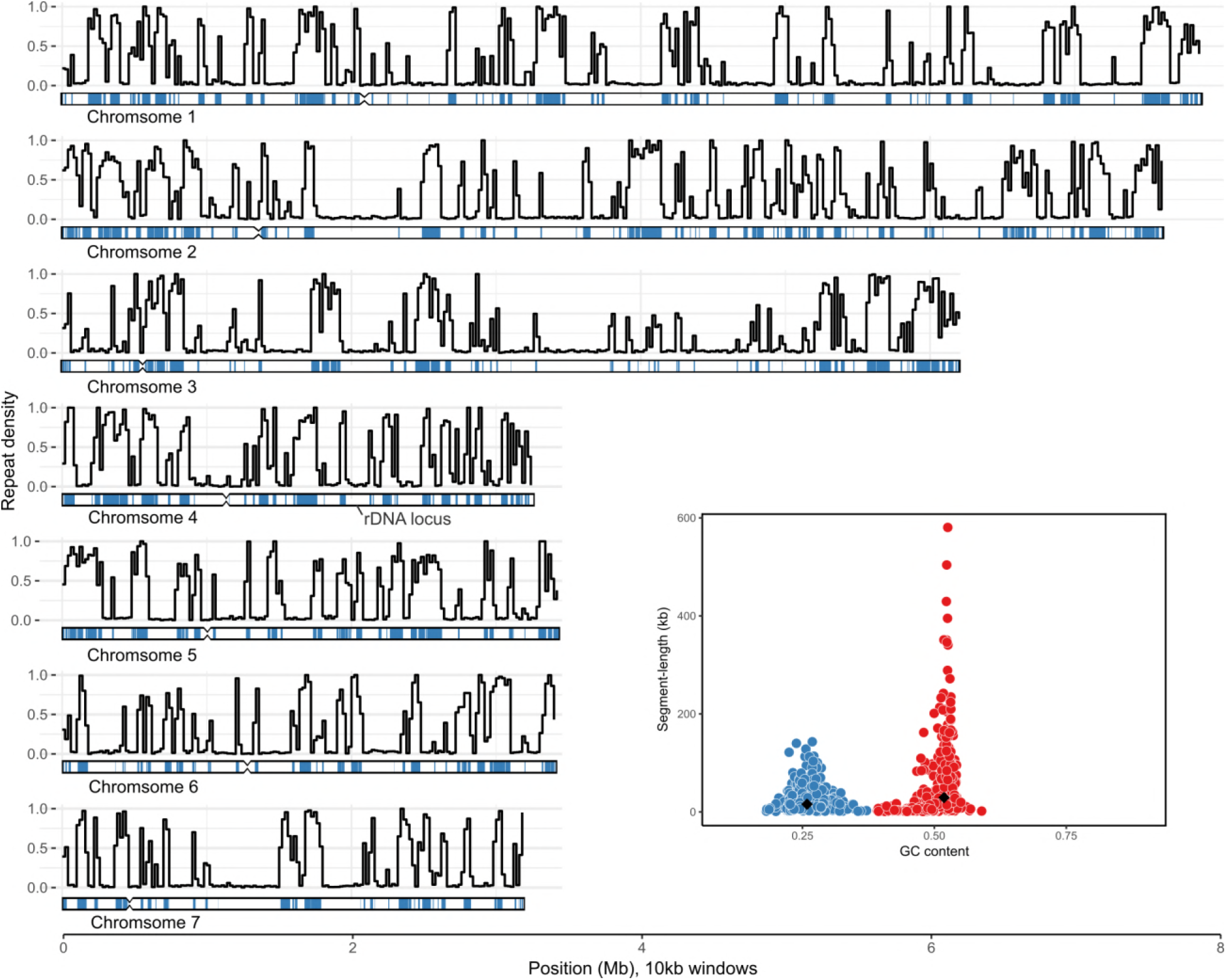
The genome comprises distinct AT-rich and GC-rich regions. Each sub-plot represents data derived from one of the seven *E. festucae* Fl1 chromosomes. The black lines represent the proportion of bases in a given 10 kb window that are part of a known repetitive element. An ideogram displaying chromosome structure is placed under each line graph. The blue tracks on each ideogram show the positions of AT-rich regions identified with OcculterCut. Notches represent the putative locations of centromeres. Inset: a scatterplot showing the length and GC-content of AT-rich segments (blue) and the rest of the genome (red); the black diamonds represent the median values for each measurement for each type of region.

**Figure 3:**
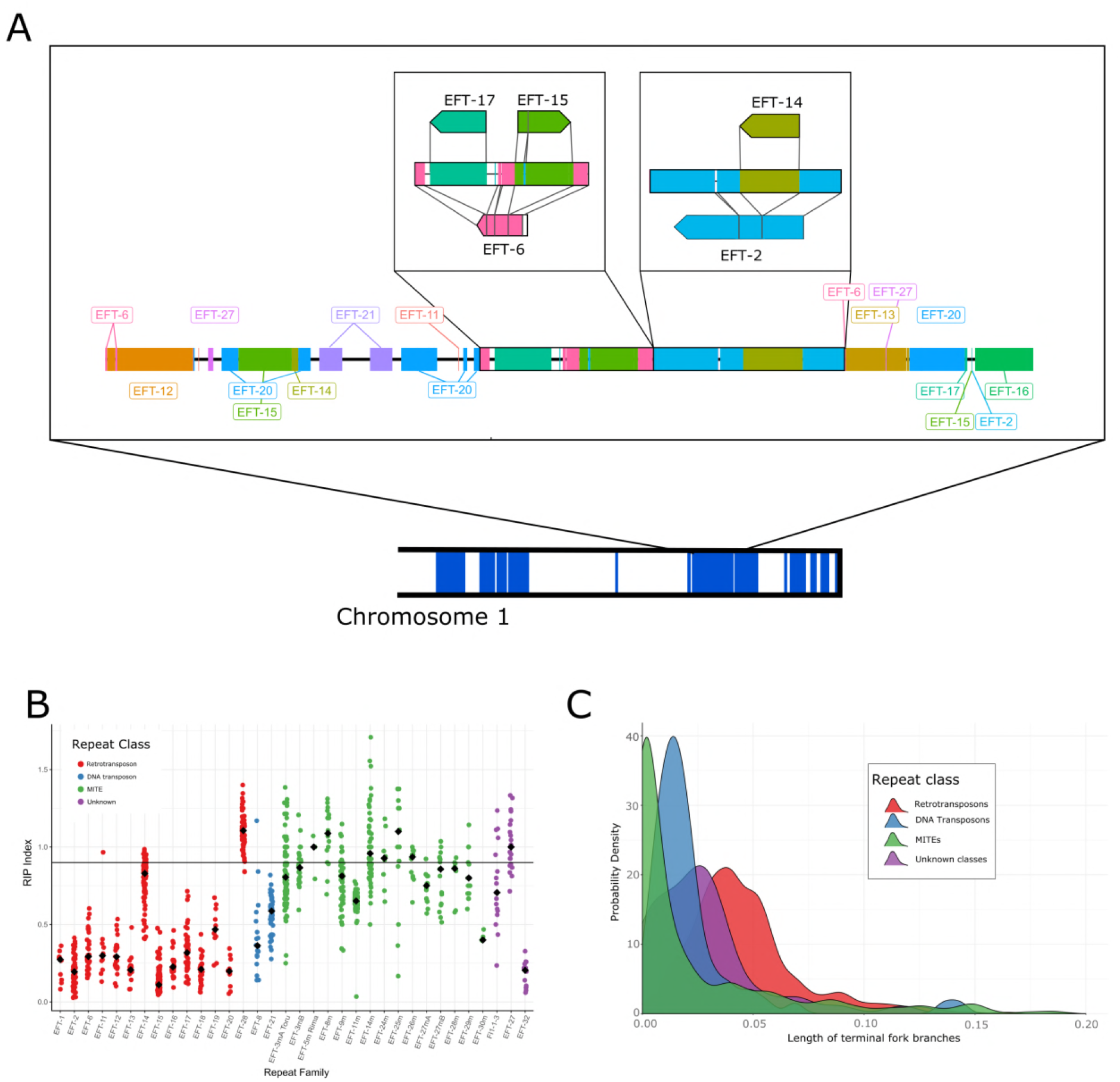
Repetitive elements have largely been inactivated by RIP and interruption by the insertion of other elements. **A**. Detailed annotation of a representative AT-rich region on the long arm of Chromosome 1. Repeats are shown as tracks along a chromosome and are color-coded by family. The two insets illustrate the nested arrangement of repeats. The reference repeats are shown above and below the genomic organization, and illustrate how the genomic repeats align to their respective reference sequences. As an example, the left-most box shows a copy of EFT-6 that has been interrupted by the integration of elements from the families EFT-17 and EFT-15. B. The RIP index, where lower values correspond to stronger evidence for RIP, was calculated for each annotated repeat. Black diamonds represent median values for each repeat family; coloured points show the full distributions of RIP index values. Points are color-coded to reflect repeat class. The horizontal line at 0.9 represents the minimum value expected for sequences free from RIP (Margolin et al. 1998). C. A phylogenetic approach was used to show the history of repetitive element invasions in the Fl1 genome. Estimated distributions of time since the last integration (measured as the number of non-RIP mutations per site on terminal branches) are plotted for all copies of each repeat class, with smaller values indicating more recent integrations.

**Table 1:** Summary information about the AT- and GC-rich components of the genome. ‘Peak GC’ refers to the mode of the nucleotide content distribution for this component, as estimated by OcculterCut. ‘Length’ is the mean length of these components in kb, with standard deviations given in parentheses. ‘Genes’ is the total number of genes (introns and UTRs included) that at least partially overlap with a given block-type (meaning genes that overlap the boundary between an AT- and GC-rich region will be counted twice).

| Component | Peak GC | Length (sd) | Total Mb | Repeat Mb | Exon Mb | Genes |
| --- | --- | --- | --- | --- | --- | --- |
| AT-rich | 25% | 25.7 (25.5) | 11.2 | 9.49 | 0.01 | 319 |
| GC-rich | 52% | 55.3 (74.7) | 23.8 | 0.2 | 13.3 | 8,844 |

### Long repeats have been inactivated by RIP

Some ascomycete fungi target repetitive elements for hypermutation using a process called repeat induced point mutation (RIP; Cambareri et al. 1989). RIP induces C-to-T mutations in large (greater than ~550 bp) sequences present as multiple copies in the genome. The action of RIP is focused on particular dinucleotide sequences, with CpA dinucleotides the most common target in most species (Cambareri et al. 1989; Braumann et al. 2008). The RIP index of Margolin et al (1998) measures the depletion of RIP dinucleotides, while accounting for the overall AT-content of a genome. Low values of this index (< 0.9) correspond to a strong signal of RIP, while larger values are expected for unaffected sequences. The long repeats that make up the AT-rich regions in the Fl1 genome have low RIP index values (Figure 3B), suggesting that the AT content of these regions is the consequence of RIP recognizing the repeats and introducing C-to-T mutations.

The presence of many copies of different repeat families, each with strong evidence for RIP, suggests a history of invasion of the genome by repetitive elements followed by their inactivation by RIP. We examined the temporal dynamics of these invasions by estimating phylogenies from the different repeat copies identified for each family. The length of terminal fork branches (i.e., branches from a single node that each lead to a tip) provide an estimate of the time at which each repeat family last integrated a copy into the Fl1 genome. To differentiate spontaneous mutations accrued over evolutionary time from the more directed RIP-induced mutations, we estimated trees from alignments in which all CpA dinucleotides were masked. When the distribution of the lengths of terminal forks is considered across classes of repeats, it is clear that the majority of retrotransposons are relatively old and have not been active in the genome for some time (Figure 3C). In contrast, DNA transposons and especially MITEs show a preponderance of short terminal branches, suggesting repeats from these families have been active more recently. The recent activity of MITEs is consistent with them avoiding RIP, potentially through their small size, although they are thought to require a transposase from elsewhere to move (Feschotte et al. 2002). These results are also consistent with the retrotransposons that have a low RIP index being inactive, as these show no evidence for recent transposition activity.

We used the RIP index values to examine which repeat families may still have functional copies within the genome (Figure 3B). Most DNA transposon and retrotransposon families are only represented by copies that have been heavily affected by RIP, suggesting these repeats are no longer functional. Repeat families EFT-8, EFT-11 and EFT-14 all have some copies with relatively high RIP index values (> 0.9), but these sequences are all either truncated (less than 85% of full length) or contain truncating stop codons (Supplementary Table S1). The non-autonomous retrotransposon EFT-28 appears to have evaded RIP entirely, presumably as a result of its small size (reference length 531 bp). In contrast MITEs appear to be only weakly affected by RIP, most likely due to their short length. There are a number of repeat families in the Fl1 genome that are not currently classified as belonging to a known repeat class. These unclassified repeats include two shorter elements (Fl1-1-3 and EFT-27, reference lengths < 610 bp) that show relatively little evidence for RIP, as well as one longer element (EFT-32, reference length 2877 bp) that shows a strong RIP signal. Thus, there are likely few or no functional autonomous transposons present in the Fl1 genome but MITES and unclassified repeats probably continue to propagate.

### Repeat rich regions structure the genome in three dimensions

The HiC data provide information on the frequency with which regions of the genome interact with each other in the nucleus, and thus provide insight into the three-dimensional structure of the genome. Contact maps generated from the HiC data reveal that three-dimensional genome structure is influenced by the AT- and gene-rich blocks. Contacts between blocks of the same type are enriched, while there are relatively few contacts between the AT- and gene-rich blocks (Figure 4A). We quantified this effect globally by using principal component analysis to extract the dominant signal from the contact data for each chromosome. The first principal component (PC) differentiates the AT- and gene-rich blocks of the genome, such that AT-rich blocks generally have high scores for this principal component (Figure 4B). Contacts between chromosomes are also dominated by interactions between blocks of the same type. This effect is strongest for AT-rich blocks, with contacts between such regions accounting for almost half of all inter-chromosomal contacts, despite these blocks comprising only a third of the genome (Figure 4C; Supplementary Figure S4). These results suggest that the AT-rich blocks are key determinants of 3D genome structure, and that the genome is partitioned into AT-rich and gene-rich blocks in three dimensions as well as in one dimension.

**Figure 4:**
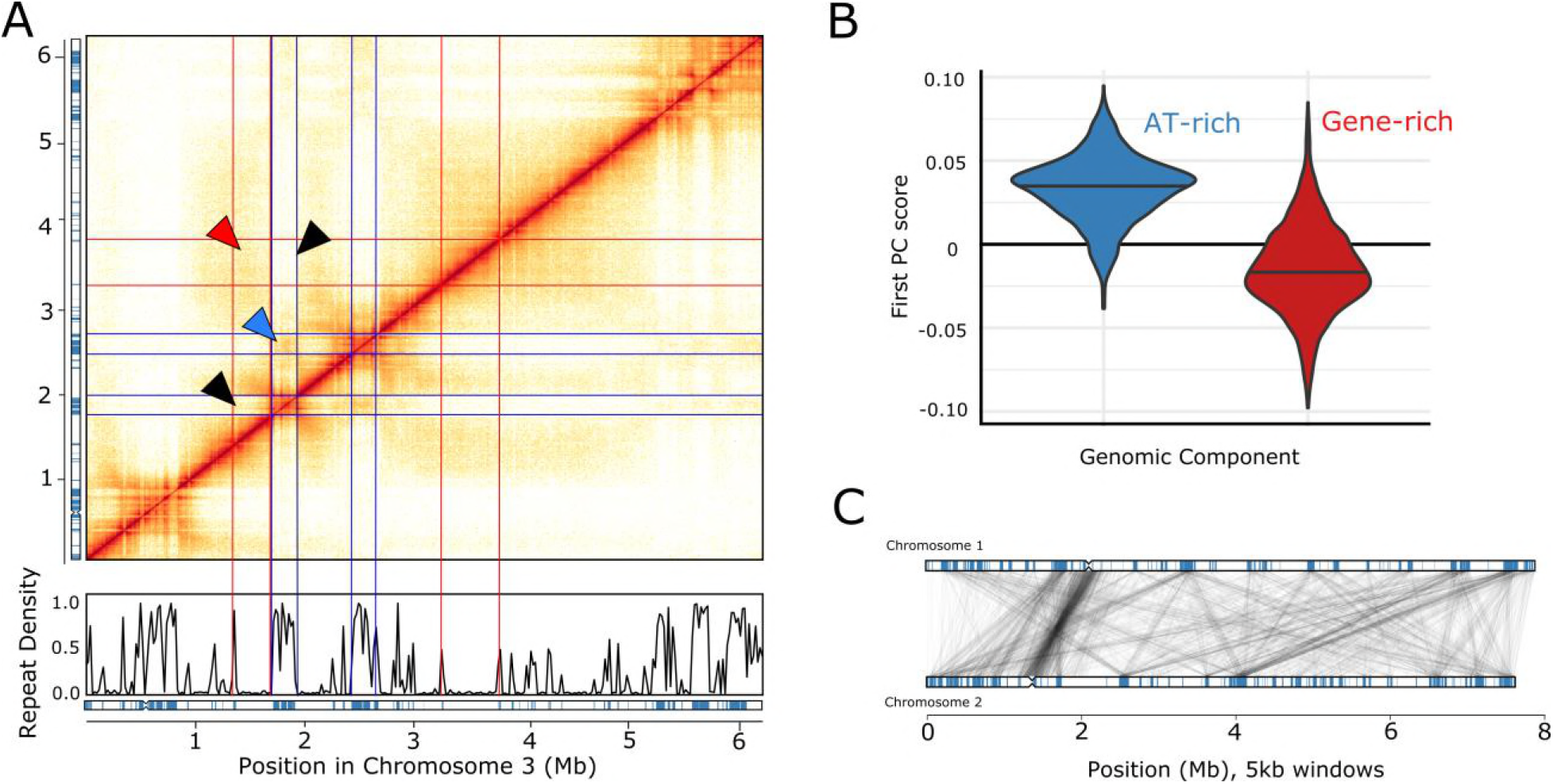
HiC data reveals interactions within and among chromosomes. **A**. Each element of the matrix reflects the frequency of contacts between two genomic windows in an exemplar region of chromosome 3. Repeat density and the locations of AT-rich regions (blue) are plotted below the matrix, as in Figure 2. Red lines represent the boundaries of gene-rich regions, blue lines boundaries of AT-rich regions. Triangles highlight examples of interactions among specific genomic regions. Red, an interaction among gene-rich regions with high contact frequency (dense shading); blue, an AT-AT interaction with high contact-frequency (dense shading); black, interactions with low contact frequency between gene-rich and AT-rich regions (light shading). B. Distribution of first principal component scores estimated from HiC data for 5 kb regions entirely made up of AT-rich (blue) or gene-rich sequence (red). C. Inter-chromosomal contacts between chromosomes 1 and 2 are shown. All 5 kb windows sharing more than five HiC contacts are connected by a grey line. The AT-rich blocks in each chromosome are indicated (blue boxes).

The AT-rich component of the genome is greatly enriched for repeats, and it might therefore be argued that the evidence for AT-AT interactions in our data might be an artefact of incorrectly mapped reads from repetitive elements. However, >97% of the genome (including 92% of the AT-rich sequences) are uniquely mappable with 80 bp reads, and the analyses discussed above include only reads that uniquely map to one location in the genome without any mismatches. Further, if the apparent enrichment of contacts between two AT-rich sequences were a consequence of incorrectly mapped reads, we would expect repeat blocks that share more HiC read pairs to contain repeats (and hence sequences) from the same family. However, we found no correlation between the repeat families found in AT-rich regions and the number of HiC contacts observed for those regions (Mantel test of correlation between distance matrices estimated from HiC contacts and shared repeat families, *r* = −2 × 10^−5^, *P* = 0.92). Therefore, we conclude that the interactions we observe between AT-rich blocks are likely to be biologically relevant.

Topologically associated domains (TADs; Nora et al. 2012) are contiguous regions of a single chromosome that have a high rate of self-interaction. In a common theme, the TADs correlate strongly with the patchwork structure of the genomic components (Supplementary Figure S5) Most AT-rich blocks contain a single TAD (mean = 1.1, sd = 0.4), while gene-rich blocks often contain multiple TADs (mean = 2.3, sd = 2.0). TADs within gene-rich blocks have a mean length of 0.14 Mb (sd = 0.19 Mb), similar to those estimated from yeasts (Eser et al. 2017). In animals, TAD boundaries are determined in part by the protein CTCF binding to particular sequence motifs (Hou et al. 2008; Dixon et al. 2012). The *E. festucae* Fl1 genome does not contain a homolog for CTCF, and we were not able to identify any sequence motifs associated with TAD boundaries, suggesting another mechanism is responsible for forming TAD-boundaries in *E. festucae*.

The HiC data also allow us to estimate the local chromatin state for different parts of the genome. The probability of observing HiC contacts between two regions of the same chromosome is expected to scale following a power law with regards to the distance between those regions in the genome sequence (Nicodemi & Pombo 2014). The parameters of this power law relationship depend on local chromatin state. When chromatin is condensed, genomic regions that are distant from each other in the one-dimensional genome are brought together, meaning the rate of interaction decays relatively slowly with respect to genomic-distance. In contrast, open chromatin states lead to relatively rapid decay rates.

Fitting these models to the Fl1 data shows that the power law exponent α, and thus the rate with which interactions decay, differs greatly between the AT-rich and gene-rich blocks. AT-rich have a relatively low decay rate (α = −0.55), while interaction frequency decays rapidly in gene-rich blocks (α = −1.29). These results suggest the AT-rich blocks are generally condensed (at least in the culture conditions assayed here), while gene-rich blocks have a more open chromatin state.

### Genome structure is associated with the regulation of gene expression

TADs have been shown to mediate the regulation of gene expression (Ulianov et al. 2015; Andrey et al. 2013; Gonzalez-Sandoval & Gasser 2016). We assessed the contribution of TADs to the regulation of gene expression in *Epichloë* by analysing RNAseq data generated *in planta* and in culture. We tested for co-regulation of genes within TADs by fitting a general linear model in which log_2_ fold difference in gene expression between plant and culture for a given gene is predicted by TAD membership. This model provides a substantially better fit than a null model (ΔAIC = 512.2), suggesting genes within a given TAD do tend to have similar gene expression changes. This model also allows us to identify the TADs with the strongest signal of co-regulation among genes. Interestingly, TADs with the strongest evidence for shared expression changes (Figure 5A, Supplementary Table S2) are enriched within subtelomeric regions of the chromosomes.

**Figure 5:**
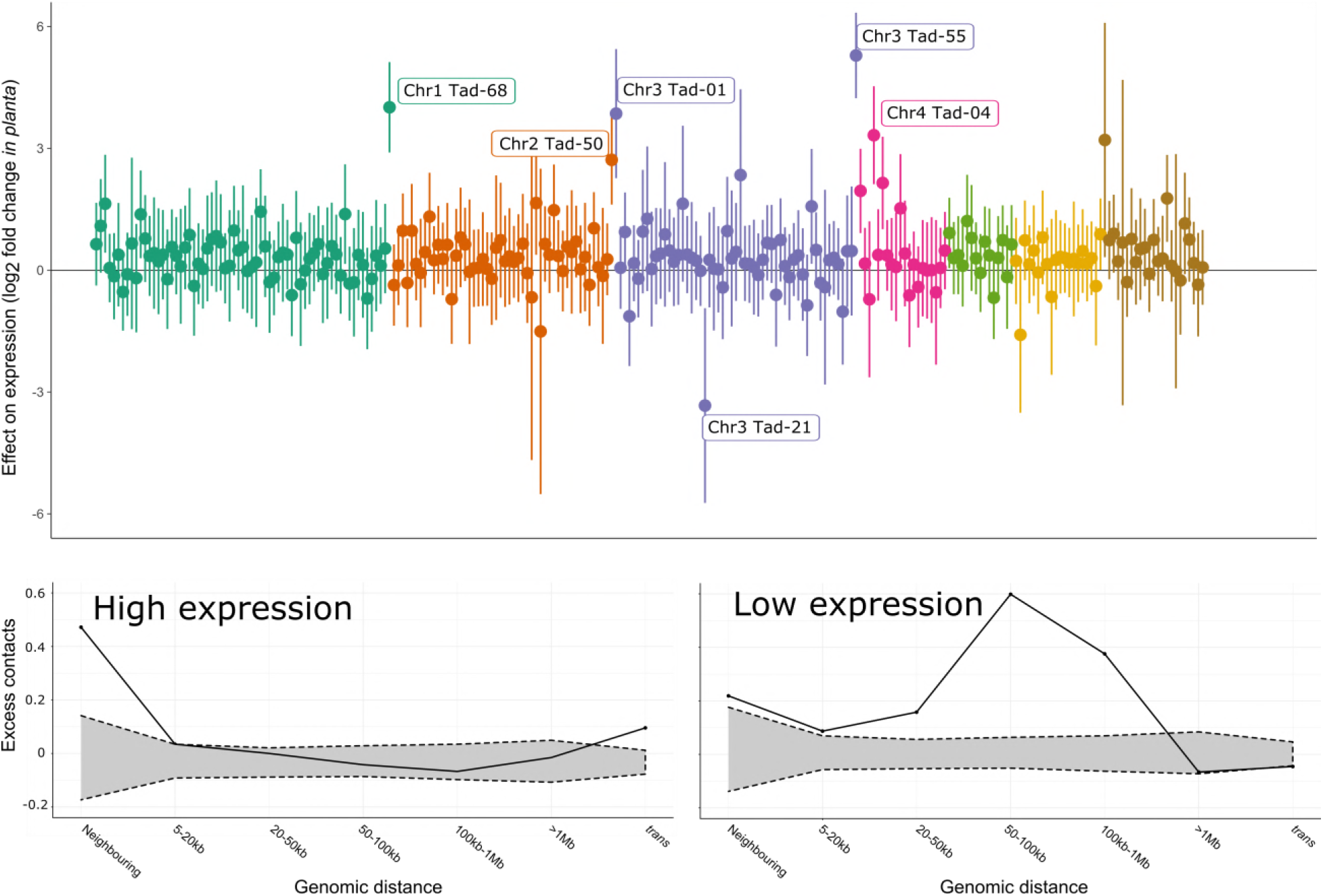
The three-dimensional structure of the genome is associated with regulation of gene expression. A: Each point is a TAD; the estimated log_2_ fold difference in expression between *in planta* and in culture conditions for all genes within that TAD are plotted. Vertical lines represent uncorrected 95% confidence intervals of this estimate; TADs are coloured by chromosome. TADs with statistically significant effects after applying a false discovery rate of 0.01 are labelled by chromosome and TAD number. Genes contained in these TADs are listed in Supplementary Table S2. B: The proportional excess ([observed – expected]/ observed) of contacts between members of the 5% of genes with the highest expression in culture. Along the x-axis, these interactions are broken down by the genomic distance between interacting gene pairs. 95% confidence intervals for expected number of contacts (grey shaded area) for each observation were obtained by counting the number of contacts from 1,000 random gene sets. C: As with B, but for the 5% of genes with the lowest expression in culture.

To illustrate how the chromatin states and three-dimensional structures identified above may contribute to co-regulation of genes, we consider one well-characterized subtelomeric gene cluster. The *EAS* cluster, which encodes proteins that catalyse the production of bioprotective ergot alkaloids, forms a part of a single TAD with a boundary corresponding to the end of an AT-rich block in culture (Figure 6). The majority of *EAS* genes have much higher expression *in planta* than they do in culture, while the expression of genes in the neighbouring TAD is largely unaffected. The relatively low expression of the *EAS* genes in culture and the fact these genes form a self-interacting domain that includes AT-rich blocks may be explained by the condensed chromatin state generally associated with AT-rich blocks extending across the entirety of this TAD. If so, the massive increase in expression of these genes *in planta* may be the result of restructuring of this region of the genome to open the chromatin up to allow gene expression. However, performing an *in planta* HiC experiment to test this hypothesis is not currently technically feasible, as DNA from the relatively small *Epichloë* genome is swamped by the more common and much larger host genome.

**Figure 6:**
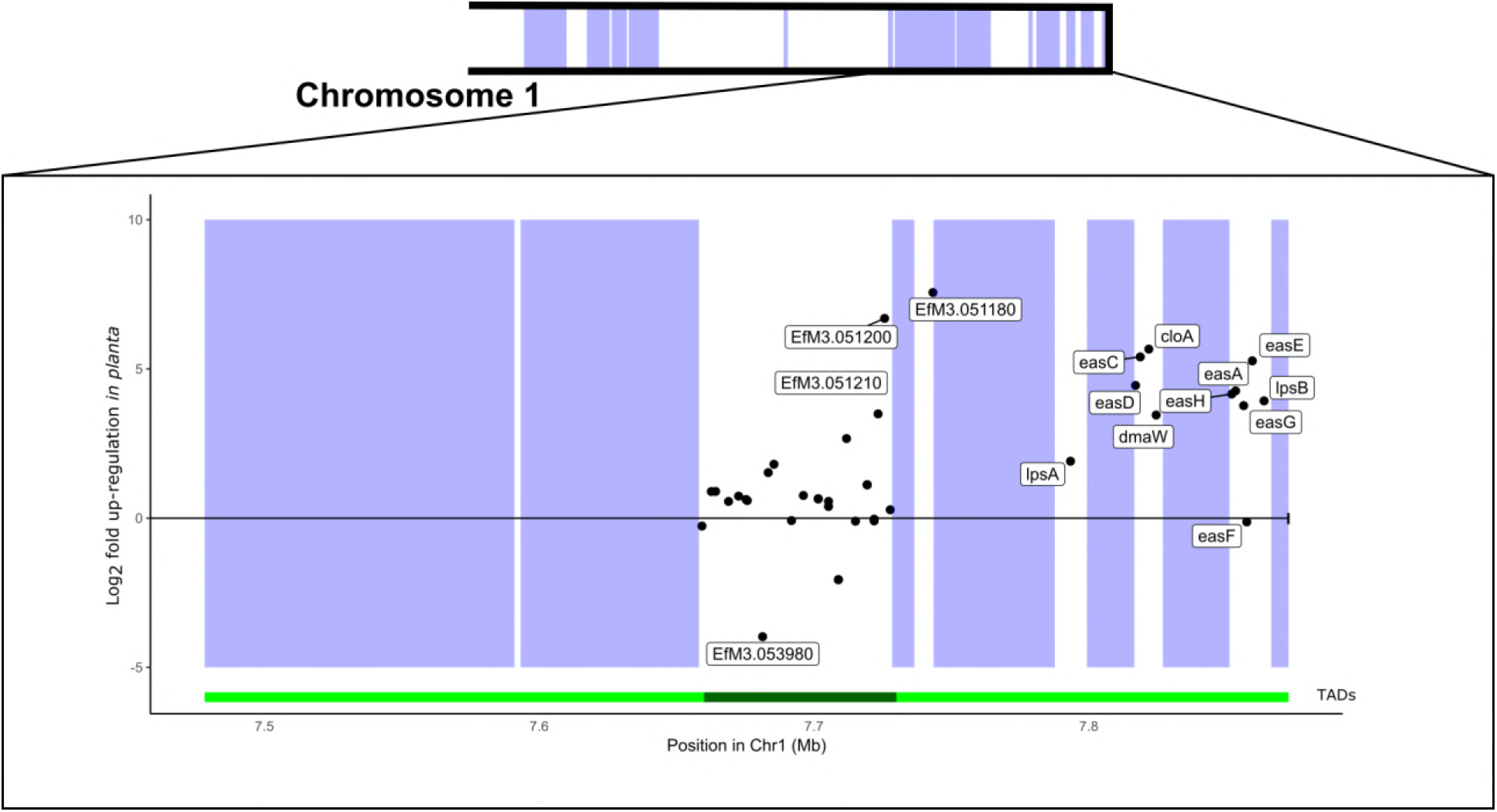
The *EAS* cluster is part of a co-regulated TAD whose boundaries are determined by repeat content. A schematic of the long arm of chromosome 1 is shown on top. Below, the genomic location (x-axis) and difference in gene expression *in planta* v culture (y-axis) for each gene is represented by a point. Genes with >8-fold difference in gene expression are labelled, as are all genes in the *EAS* cluster. Blue boxes indicate AT-rich blocks. The green tracks below the graph represent the location of distinct TADs.

To further explore the role of nuclear architecture on gene expression, we used the HiC data to test whether genes with similar expression levels tend to have more frequent interactions, suggesting localization to similar nuclear positions. We find that highly expressed genes (the 5% of genes with highest expression in culture) tend to interact with each other more than expected by chance. These interactions are most over-represented among genes that are either immediately adjacent to each other or are located on different chromosomes. In contrast, interactions between highly expressed genes are relatively uncommon between genes located at intermediate distances within a chromosome Figure 5B). The 5% of genes with the lowest expression in culture also tend to interact with each other more commonly than would be expected by chance. In contrast with the highly-expressed genes, these interactions are most overrepresented at intermediate genomic scales (Figure 5C).

### Repeats are associated with changes in gene expression

Repetitive elements have repeatedly been shown to provide novel genetic material that can be co-opted by host genomes (Chuong et al. 2017). For fungi in particular, it has been suggested that the presence of repeats can create a “two-speed genome” in which repeat-rich regions of the genome evolve more quickly and therefore provide a source of genetic novelty that natural selection can act on (Dong et al. 2015; Faino et al. 2016). We examined the possibility that the AT-rich blocks contribute to adaptive evolution in *Epichloë* by testing for an association between these blocks and genes likely to be involved in host interactions or which may underlie lineage-specific responses (Figure 7). Genes with apparent Fl1-specific expression (defined here as genes that are expressed *in planta* by strain Fl1 but do not have an orthologous gene model in the current reference strain E2368) are more than twice as likely to fall within 5 kb of an AT-rich block compared to other genes (odds ratio = 2.32, 95% CI = [1.82, 2.95]). Genes that are differentially expressed *in planta* are also significantly over-represented near these blocks (odds ratio = 1.62, 95% CI = [1.31, 2.01]). However, there was no significant association for genes that encode secreted proteins (odds ratio = 0.85, 95% CI = [0.66, 1.09]) or genes involved in the production of secondary metabolites (odds ratio = 1.27, 95% CI = [0.89, 1.89]).

**Figure 7:**
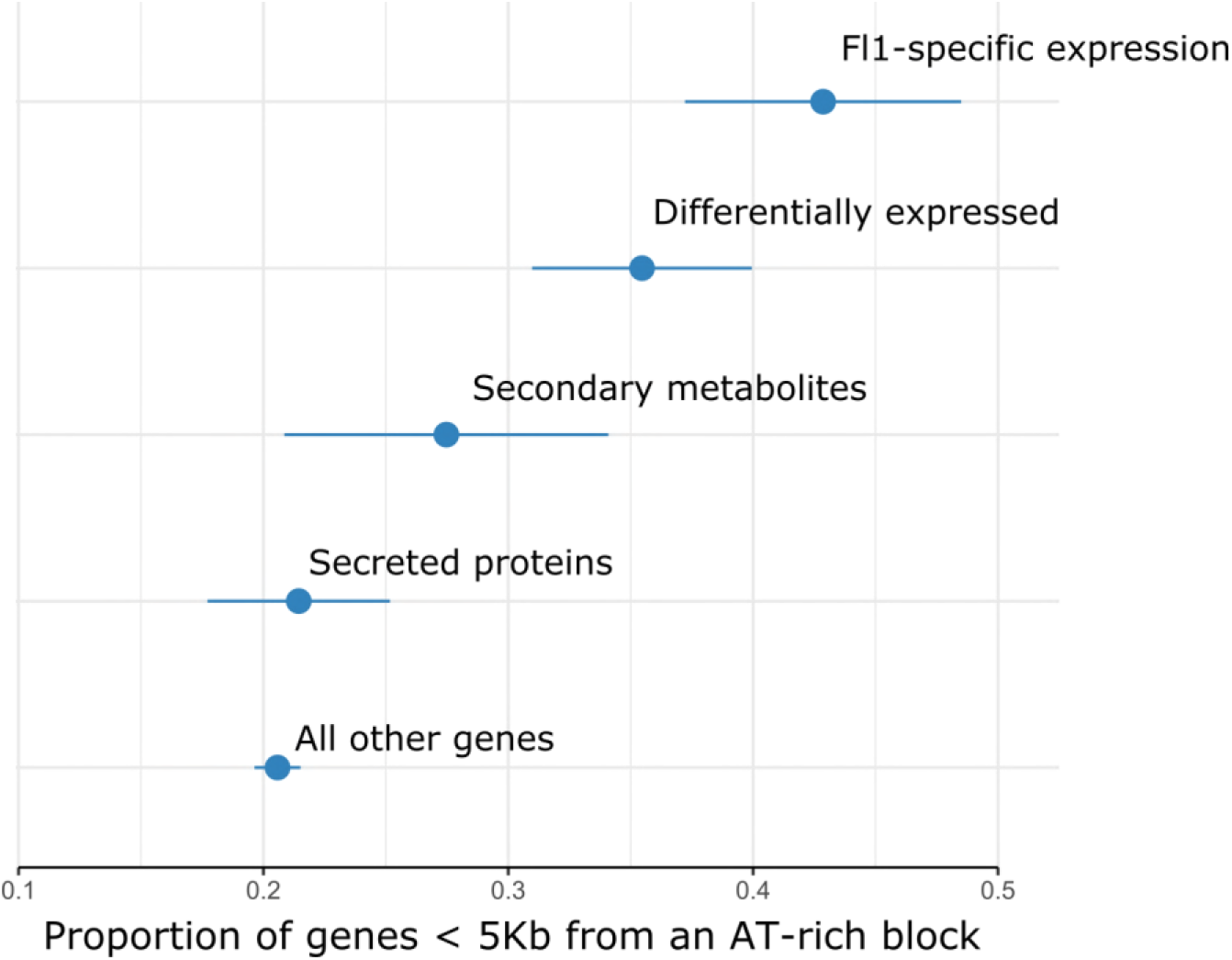
Differentially expressed genes and genes with Fl1-specific expression are more likely to occur near AT-rich blocks. We identified four classes of genes that we expect to play important-roles in host-adaptation in *Epichloë*. Each point represents the proportion of genes in a given functional class that fall within 5 kb of an AT-rich block. ‘Secreted proteins’ = genes that encoded proteins with signal peptides, ‘Differentially expressed’ = the 5% of genes with the highest fold-difference increase in gene expression *in planta* compared to culture, ‘Fl1-specific’ = genes with no evidence for expression in the reference strain E2368, and ‘secondary metabolite’ = genes known to be involved in production of secondary metabolites including terpenes, indole diterpenes and ergot alkaloids (see Methods for details).

Although gene-rich regions of the genome are almost free from long repeats, they do contain a number of the smaller MITE elements. These elements are known to act as regulators of gene expression both plants and plant-associated fungi, so we tested whether MITES were more likely to appear near genes that are differentially expressed *in planta*. We find the 20% of genes with the highest differential expression *in planta* are almost three times more likely to have a MITE within 2 kb of their coding sequences (14.2% of such genes, 5.2% of all others). The over-representation of MITEs near up-regulated genes is not uniform among MITE families; some show almost no effect, while others such as EFT-9m are more than five-fold overrepresented (Figure 8A). The over-representation of MITEs is strongest for the genes that show the greatest differential expression *in planta*, where the signal appears immediately upstream of the transcription start site (Figure 8b).

**Figure 8:**
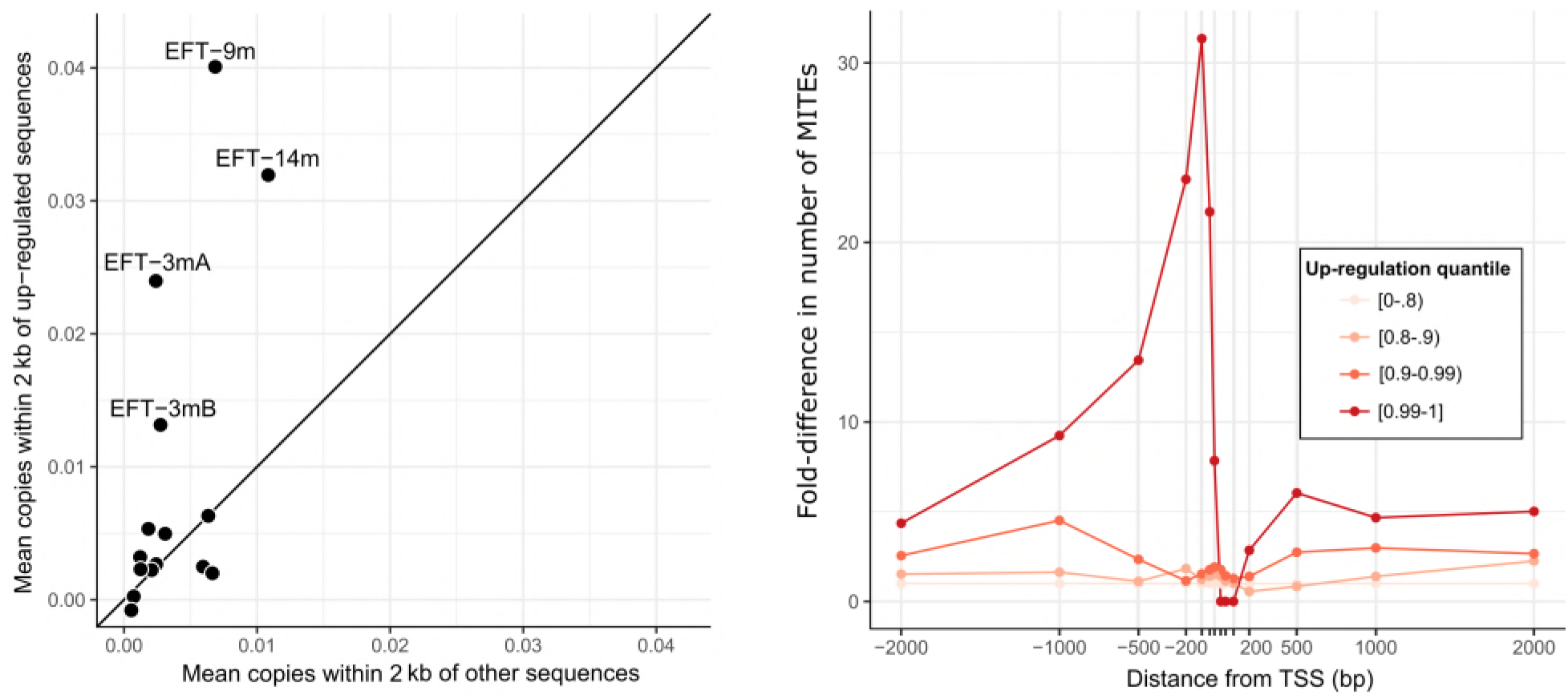
Some MITE families are over-represented near highly differentially expressed genes. A: Each point represents the mean number of copies of a given MITE family within 2 kb of genes that are differentially expressed *in planta* (x-axis) vs the same measure for all other genes (y-axis). Families with more copies near differentially expressed genes fall above the diagonal line. Those families with the greatest over-representation are labelled. B: Fold-difference in the number of MITEs near differentially expressed genes plotted by position relative to the transcription start site (TSS; negative values correspond to upstream positions). Four differential gene expression quantiles are plotted, with darker red points corresponding to genes with the greatest differences in expression.

### m6A does not appear to mark gene expression

The PacBio sequencing data also contain information about chemically modified bases. The only eukaryotic modification that could be reliably diagnosed from this data is N^6^-methyladenine (6mA). An association between 6mA marks and gene expression levels has recently been suggested (Mondo et al. 2017). Approximately 0.04% of adenines show evidence for this modification, but the modified bases are approximately evenly distributed between the AT-rich (0.042% of adenines) and gene-rich blocks (0.038% of adenines), and are not associated with genes with very high or low expression. Adenines in ApG dinucleotides account for more than 50% of the m6A we detect, but again this proportion does not differ substantially between gene-rich and AT-rich blocks, or protein-coding sequences and the rest of the genome (Supplementary Figure S6). Therefore, we find no strong evidence to suggest a role for 6mA in the regulation of expression in *Epichloë*.

## Discussion

### A finished reference genome for genus *Epichloë*

Here we present a complete genome sequence for *E. festucae* strain Fl1. This chromosome-level genome was generated using PacBio, HiC, and Illumina sequencing approaches, and is a substantial improvement on the previous, fragmentary *Epichloë* genomes. It runs telomere-to-telomere for seven nuclear chromosomes, places the highly repetitive ribosomal RNA gene repeats, provides putative positions for all seven centromeres, and includes the mitochondrial DNA genome. We find a small number of bases (approximately 0.04% of adenines) with methylated adenine (6mA), preferentially on ApG dinucleotides, but find no evidence for functional enrichment of this mark. Previous genomic work in *Epichloë* has suggested a sizable repeat composition. Our complete genome illuminates not only the extent of repeat content, but also the highly patchwork nature with which it is organized. The Fl1 genome is divided into blocks of highly AT-biased, repeat-rich DNA that almost entirely lacks genes, interspersed with gene-rich blocks that have balanced GC content. While the highly repetitive nature of the AT-rich blocks is likely the reason that short read-based *Epichloë* genome assemblies have been fragmentary, the block-like structure of the genome we reveal here suggests that such short-read assemblies are likely to have captured a great majority of the coding parts of the genome within the gene-rich blocks.

### AT-rich regions: graveyards for repetitive elements and nurseries for lineage specific genes?

We find that the AT-rich blocks are most likely the result of RIP mutating the repeats to inactivate them, and thus driving dramatic increases in repeat AT content. This is consistent with most of the transposon repeats in the genome lacking a signature of recent movement, suggesting that RIP has been effective in disabling most full-length, active transposable element repeats. The AT-rich regions can thus be seen as ‘graveyards’ containing the remains of ancient transposons that have been inactivated by RIP or interrupted by the subsequent integration of other elements. In contrast, small non-autonomous MITE repeats do show evidence for recent movement, are generally not subject to RIP (presumably due to their small size which avoids the RIP mechanism), and are frequently found in the gene-rich blocks. It remains an open question as to which elements have catalysed the recent mobilization of these MITEs in the Fl1 genome, though some DNA transposons and elements not currently classified show evidence for recent mobilizations. Complete genomes for other *Epichloë* species will help to reconstruct the evolutionary history of transposable element invasions in this genus.

The patchwork nature of the Fl1 genome raises the question of what is responsible for this dichotomous ‘blocky’ pattern of genome structuring. We have shown that the AT-rich blocks contain series of nested repeats, suggesting repeated transposition of repeats into existing repeats. One possible explanation for this pattern is that transposition events into the regulatory and coding regions of the genome, particularly those of large transposons that are likely to be disruptive, are subject to strong negative selection. Therefore, we predominantly see transposons in non-functional parts of the genome (i.e., existing transposons), as only these insertion sites are tolerated. Alternatively, the preponderance of nested repeats in the genome could be a direct consequence of insertion-site preferences for transposons. Many LTR retrotransposons preferentially integrate into short AT-rich sequence motifs (Wei et al. 2013; Crénès et al. 2010). If *Epichloë* transposons have similar biases, transposition events would occur more frequently in regions previously subject to RIP. RIP could thus have a potential double-hit benefit: not only does it inactivate transposons that have invaded the genome, but when subsequent transposons invade, they preferentially insert into regions that are unlikely to perturb genome function (i.e., the AT-rich, RIP-inactivated transposons). This “safe-haven” hypothesis for nested transposition events could be tested by experimentally determining whether *Epichloë* transposons show such an AT-rich insertion bias.

Although the AT-rich regions are very gene poor, the genes they do contain may be of particular importance for strain-specific host interactions. We identified 317 genes with evidence for expression *in planta* in strain Fl1 and no ortholog in the existing transcriptome assembly for strain E2368. These genes with apparently Fl1-specific expression are greatly over-represented within AT-rich regions of the genome, suggesting these regions may indeed contribute to lineage-specific evolution. It should be noted, however, that transcriptomic data from strain E2368 was not analysed with the benefit of a complete reference genome. Though these results suggest the AT-rich regions created by repetitive elements and RIP maybe host lineage-specific genes, further work will be required to establish the degree of among-strain diversity in gene expression and its molecular basis. The first complete reference genome we report here will be an important resource for this work.

### Repeats have a profound influence on the three-dimensional structure of the genome

Within a nucleus, the three-dimensional structure of a genome is organized in a hierarchical fashion (Bonev & Cavalli 2016). Individual nucleosomes aggregate to form chromatin fibres, those fibres fold to form contiguous chromatin loops, different loops interact to form self-interacting domains within chromosomes, and chromosomes themselves take on characteristic sub-nuclear territories. In *E. festucae* Fl1 we find the repeat-rich AT-rich blocks have a profound influence on the organization of the genome. The AT-rich blocks generate more tightly packed chromatin than the gene-rich blocks. These tightly-packed blocks interact with each other more than they do with gene-rich regions, suggesting they co-locate within the nucleus and form the core of the three-dimensional structure of the genome. The repeats also structure the genome at higher levels. Moreover, the strongest long-range intra-chromosomal contact signals arise from interactions between blocks (either gene- or AT-rich) of the same kind. Similarly, the majority of inter-chromosomal interactions occur between blocks of the same kind. Taken together, these results show that the interleaved AT- and gene-rich blocks shape the genome in ways that bring these elements together in three-dimensional nuclear space to the exclusion of the opposite block type.

In many species, the boundaries of TADs are defined by proteins that bind to particular sequence motifs. We were not able to find any sequence motifs associated with the boundaries of TADs in the *E. festucae* genome. The patchwork structure of this genome may partially obviate the need for sequence motifs to recruit proteins that define TAD boundaries. There is evidence that the molecular machinery associated with RIP can detect repetitive elements and mark them with methylated cytosines, which, in turn, could drive the formation and spread of condensed chromatin around these repeat regions (Gladyshev 2017; Gladyshev & Kleckner 2017). Under this model, the condensed chromatin states formed by the AT-rich blocks would act to insulate the relatively open chromatin loops of the gene-rich blocks from each other, thus naturally creating TAD-like structures in the genome (Figure 9).

**Figure 9:**
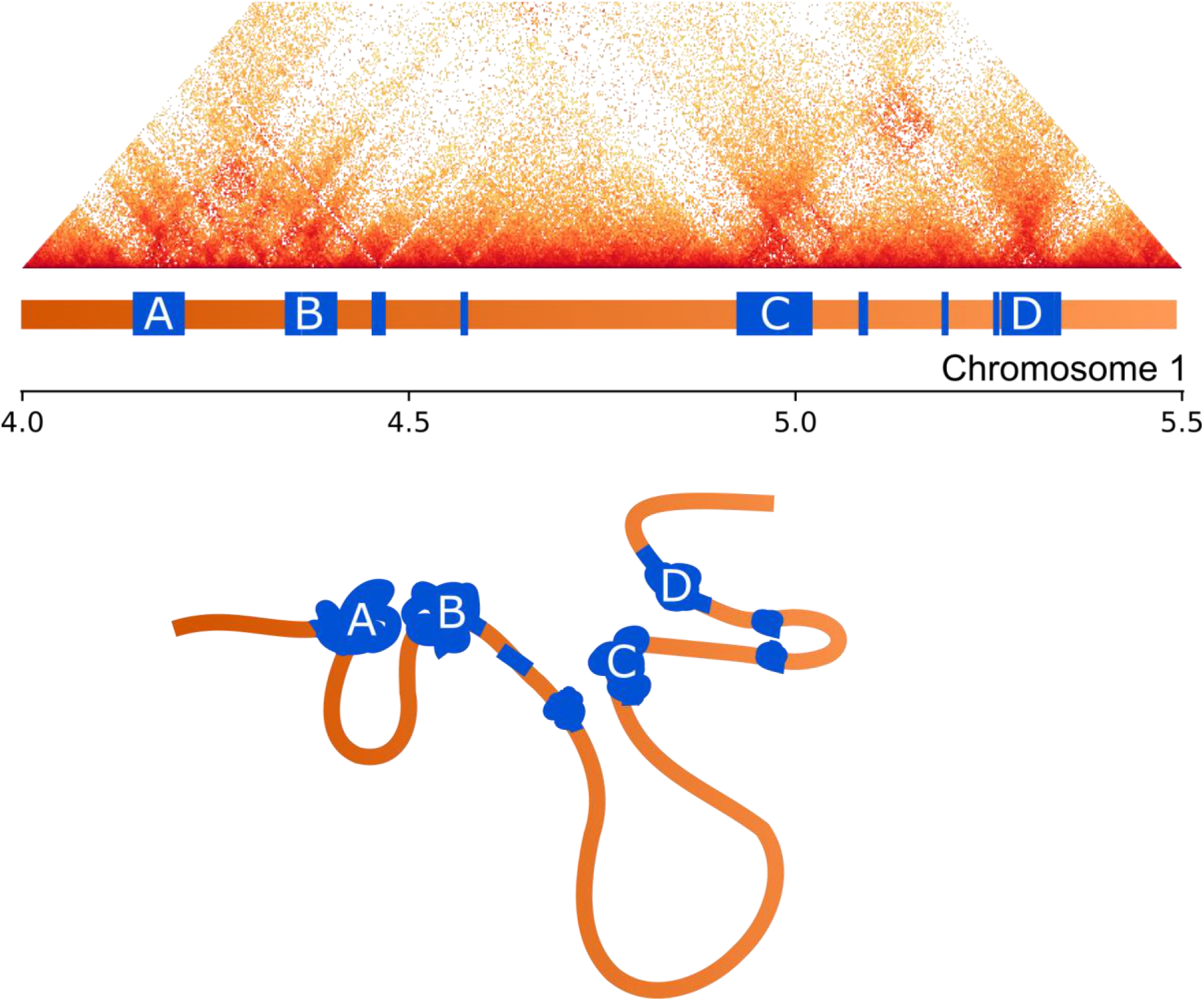
Influence of repeats on chromatin folding. Above: HiC interaction data for an exemplar region of chromosome 1 displayed as a triangular matrix with darker shading corresponding to higher interaction frequency. Directly below his matrix the structure of the 1-D genome is shown with gene-rich regions shaded orange and AT-rich regions blue, the largest AT-rich regions are labelled A-D. Below: schematic showing an interpretation of the contact data for this region. Shading and labels follow from above. The repeat regions are condensed and pulled together to form regions with high interaction frequencies (corresponding to darker colours in the HiC data). Gene-rich regions form open chromatin loops, with varying levels of chromatin packing, which are isolated from each other by the interaction of condensed AT-rich blocks.

Nevertheless, larger gene-rich blocks frequently form more than one TAD, suggesting that additional TAD formation mechanisms are also likely to be in operation. For instance, in *Neurospora crassa*, another filamentous fungus, chromatin states are generally associated with particular histone modifications (Klocko et al. 2016). The reference genome we report here will be an important resource for future studies that use epigenetic modification, chromatin accessibility and immunoprecipitation assays to uncover the molecular mechanisms by which these three dimensional structures are formed and maintained.

### Genome architecture influences environment-specific gene expression

Establishing a symbiotic relationship infection of the host grass requires major transcriptional reprogramming in *E. festucae*, which is in part achieved by extensive chromatin remodelling (Chujo & Scott 2014). These changes allow the fungus to interact with its host and produce the secondary metabolites that protect the host from pests and environmental challenges. Previous studies have been able to establish which genes are differentially expressed *in planta* versus in culture, and in mutants that are unable to establish productive symbioses (Eaton et al. 2010, 2015). Using our complete genome sequence and HiC data, we can examine the regulation of these genes in a broader spatial genomic context.

Interest in the AT-rich regions present in the genomes of some filamentous fungi has largely focused on their evolutionary importance. We suggest these regions may also play an important role in regulating gene expression. Genes that are differentially expressed *in planta* compared to culture are more likely to fall near an AT-rich region than other genes. The *EAS* cluster, a sub-telomeric gene suite that encodes proteins responsible for bioprotective ergot alkaloid production (Fleetwood et al. 2007; Schardl et al. 2013), provides an illustrative example of how chromatin remodelling around the AT-rich regions may contribute to gene regulation. In culture, the *EAS* cluster forms a single TAD with a boundary that corresponds to the edge of an AT-rich region. Genes within this TAD are much more strongly expressed *in planta* than in culture, while genes in a neighbouring TAD appear to be largely unaffected. We suggest this pattern is a result of the AT-rich regions associated with the *EAS* cluster forming a condensed chromatin state in culture (as do most AT-rich regions) and chromatin remodelling *in planta* being key to regulation of these traits. We show that *in planta* differentially expressed genes, including many genes important for the *Epichloë* symbiosis, are significantly more likely to be located next to AT-rich blocks. Thus, the wealth of unannotated genes near these blocks that are located in TADs with significant co-regulation should be a priority for future research.

The sub-nuclear locations of genes can contribute to gene expression, with genes having particularly high or low expression shown to co-locate in *Schizosaccharomyces pombe* (Grand et al. 2014). We find evidence for spatial regulation in *E. festucae*. The genes with the highest and lowest expression in culture (the condition from which the HiC data was generated) both interact with each other more than would be expected by chance. Strikingly, however, the distance between interacting and co-regulated genes differs between the highly and lowly expressed genes. We find that lowly-expressed genes tend to interact with other lowly-expressed genes located at intermediate genomic distances (up to 100 kb apart), which we suggest reflects the association of condensed chromatin state acting to bring genes located apart in the linear genome together in three-dimensional space. In contrast, highly expressed genes have an excess of contacts at a local scale as well as between chromosomes. This suggests that neighbouring genes are co-regulated, and that spatially distant genes are brought together to form transcription factories (Edelman & Fraser 2012).

### Repeats may directly regulate gene expression

Although most repeats in the *E. festucae* genome are restricted to the gene-poor AT-rich regions, sequences from the MITE class of small DNA repeats are common in genic regions. MITEs have previously been proposed as regulators of gene expression in *Epichloë* (Fleetwood et al. 2011) and other filamentous fungi (Schmidt et al. 2013). We show that MITEs have been transpositionally active in the recent history of *Epichloë*, meaning it has been possible for MITE integrations to contribute to adaptation to specific hosts or environments. Consistent with a direct role for MITEs in regulating gene expression, we find several MITE families that are enriched near genes with the largest differential expression between culture and *in planta* conditions. The possibility that these elements may be influencing expression of these genes is strengthened by the finding that the effect is strongest near the transcription start site of the most highly up-regulated genes and near the transcription start site of genes. We therefore suggest that gene regulatory mechanisms that recognize the sequences from specific MITE families may have evolved to enable rapid recruitment of genes into the symbiosis-expressed gene network. This would occur through selection for advantageous gene expression changes following MITE dispersion events. Similarly, it is possible that some of the other MITE families not involved in promoting *in* planta-specific expression might instead contribute to the regulation of different gene subsets during other growth conditions, such as formation of the *Epichloë* pre-sexual stromata structures. The complete genome sequence we present here will provide an important resource for future investigations into such phenomena.

## Conclusions

We have generated the first finished genome assembly for an *Epichloë* species. The assembly reveals a remarkable patchwork structure to the genome, in which large regions of very high repeat-density are interleaved with blocks of gene-rich sequence. We have shown this patchwork structure has a profound influence on the way the genome is organized in the nucleus. Importantly, this three-dimensional genome structure appears to mediate the functional organization of the genome into regions of distinct gene expression profiles. We find that repetitive regions of the genome are strongly associated with the most striking patterns of differential gene expression, and these frequently involve genes that underpin the intimate symbiotic relationships *Epichloë* species form with their host species. The reference genome we report here will be an important resource for future studies that investigate the molecular basis by which the three-dimensional structure of the genome is maintained, and identify how the regulation of particular genes, particularly those modulating symbiosis, is achieved.

## Materials and Methods

### Fungal strain and DNA extraction

*E. festucae* strain Fl1 (Tanaka et al. 2006) from the Massey University Culture Collection (accession PN2278) was grown in potato dextrose broth and harvested after four days to avoid polysaccharide build-up. Mycelia were freeze dried and 40 μg of genomic DNA was extracted using a standard phenol-chloroform-isopropanol process as per Byrd et al (1990). The DNA was further purified with the PowerClean^®^ Pro DNA Clean-Up Kit (Mo Bio, Carlsbad, CA, USA) to remove residual inhibitory compounds, yielding 20 μg of high molecular weight, high purity genomic DNA, as assessed by 0.8% agarose gel electrophoresis, spectrophotometry (Nano Spectrophotometer A260/280 and A260/230 ratios) and fluorometric analysis (Invitrogen Qubit Fluorimeter).

### DNA sequencing

Long-read PacBio sequence data were generated using 6 SMRT cells on the PacBio RS II (Sequencing and Genomic Technologies Shared Resource, Duke University, NC, USA). Short paired-end reads (2 × 250 bp) were produced from a Truseq DNA Nano library (550 bp insert size) using Illumina MiSeq technology (New Zealand Genomics Limited, Palmerston North, New Zealand).

We performed a chromosome-conformation capture experiment using high throughput sequencing (HiC). At present *in planta* HiC experiments are not feasible due to the preponderance of plant-derived sequences in DNA extractions from infected tissue, so we instead investigate the three-dimensional structure from cells grown in culture. HiC libraries were created using a modified version of the method described in Burton et al. (2014). Our approach differed in that cell pellets were ground in liquid nitrogen prior to glass beading, the restriction endonuclease Sau3AI was used to digest the cross-linked chromatin, and the KAPA Hyper Prep kit was used to create the Illumina library instead of the Illumina TruSeq kit. HiC libraries were sequenced on an Illumina NextSeq 500 (University of Washington, WA, USA), producing 80 bp paired-end reads.

### Genome assembly

We used an iterative approach to produce and then improve a genome assembly for Fl1 (Figure 1). Initially, assemblies were produced from the PacBio sequences using two different assemblers: first, with Canu v. 1.3 (Koren et al. 2017), assuming a genome size of 35 Mb (estimated from earlier partial assemblies) and allowing an error rate of 0.025; and second, with HGAP from the SMRT Analysis Suite v. 2.3.0 (http://www.pacb.com/products-and-services/analytical-software/smrt-analysis/), assuming a genome size of 35 Mb and using default parameters, followed by polishing with the Quiver algorithm from the same software suite. As both assemblies were still partially fragmented (six contigs contained telomeric repeats on only one end), HiC reads were used to identify contigs from the Canu assembly that were connected by a large number of HiC read-pairs. This produced a scaffolded assembly, which included gaps of unknown size between contigs that had been joined with the HiC data. We found the alternative algorithm used by the HGAP assembler was able to bridge the five gaps in our Canu assembly, so we used HGAP-derived sequences to fill these gaps. Finally, the manually scaffolded Canu assembly was polished with the finisher Pilon v. 1.20 (Walker et al. 2014) correcting only base-level errors, particularly single base errors and single base indels.

### Assessing genome quality

We assessed the quality of the genome assembly in two ways. First, we aligned Sanger end sequences of fosmids previously generated from *E. festucae* strain Fl1 (Schardl et al. 2013), recorded the direction in which each read aligned, and inferred the insert size of each fosmid. Finally, we compared the sequences of a set of well-annotated genes from previously published studies to our final genome sequence.

### Genome annotation and gene expression analyses

Repetitive sequences were identified with Repeatmasker v. 4.0.6 (Smit et al. 1996), using rmblast v. 2.6.0 to perform sequence searches using as input a custom library containing known repeats in *Epichloë* spp. (Fleetwood et al. 2011; Schardl et al. 2013). AT-rich regions within the genome were identified using OcculterCut v. 1.1 (Testa et al. 2016). To find sequences affected by repeat-induced point mutations (RIP; Cambareri et al., 1989), the depletion of RIP-targeted dinucleotides was measured in each OcculterCut interval using the index of Margolin et al (1998), namely (CpA+TpG)/(ApC+GpT).

To identify protein coding genes, we first inferred the locations of existing E2368 gene models (Schardl et al. 2013) on the complete genome via alignment with blastn v. 2.4.0 (Camacho et al. 2009). We then identified putatively novel transcripts using previously generated RNAseq data (BioProject PRJNA447872). RNAseq reads derived from hyphal cells in culture and *in planta* were aligned to the new genome reference using STAR v. 2.5 (Dobin et al. 2013). The cufflinks v. 2.2.1 suite (Trapnell et al. 2012; Roberts et al. 2011) was used to assemble transcripts from these alignments. Genes differentially expressed *in planta* versus in culture were identified using DESeq2 v. 1.16.1 (Love et al. 2014).

### Detection of modified bases

6mA base modifications were detected with the PacBio SMRT Analysis suite v. 2.3.0. To remove potential false positive modification calls, we removed sites with an mQV score < 20. We used Bedtools v. 2.25 (Quinlan & Hall 2010) to calculate the percentage of adenines with modifications in AT and gene-rich regions of the genome and in regions close to transcription start sites.

### Repeat analysis

We used Bedtools v. 2.25 (Quinlan & Hall 2010) to calculate the total number of bases from each repeat family that overlap with the AT-rich and relatively GC-rich regions identified by OcculterCut. A phylogenetic approach was used to examine the dynamics of transposition within each repeat family over time. In short, all near-full-length (>85% of reference length) copies of those repeat families represented by at least ten such copies in the genome were extracted with Bedtools. We produced a multiple sequence alignment for each family using MUSCLE v. 3.8.31 (Edgar 2004). To minimize the effects of RIP mutations on our phylogenetic estimates, we replaced all TpA dinucleotides (which can be produced by CpA → TpA mutations) with NN. We then estimated phylogenies from each alignment with RAxML v. 8.2.3 (Stamatakis 2006), using the GTR model of DNA substitution with rates of evolution among sites following a gamma distribution. Terminal fork branch lengths were extracted from each tree using the R package ape v.4.1 (Paradis et al. 2004).

### HiC analysis

Before aligning our HiC reads, we generated a ‘mappability mask’ for the reference genome. Artificial 80 bp reads covering every base of the reference sequence were produced and these ‘reads’ were then mapped back to the assembly. We recorded the source positions of simulated reads that could not be mapped to their correct position with confidence (i.e., reads that mapped to the wrong position, reads that could not be uniquely mapped and reads with mapping quality < 30) to produce a mask of difficult-to-map positions.

We independently aligned forward and reverse HiC reads to the complete genome using bwa mem v. 0.7.15-3 (Li 2013). Reads with a mapping quality phred score < 30, or those that were mapped to positions overlapping the poor mapping mask, were removed from alignments prior to subsequent analysis. A ‘contact matrix’ representing the number of interactions among regions of the genome at a 5 kb resolution was produced from this alignments using HiCdat v. 40f4814 (Schmid et al. 2015), making use of the iterative normalization procedure described by Zhang et al (2012). We used the R (R Core Team 2017) package associated with HiCdat to investigate properties of this contact matrix. Specifically, we fitted power law models explaining the rate of interaction among genomic regions as a function of genomic distance. These models were fitted to the complete genome, but also separately to the subset of AT- or relatively GC-rich regions that are >100 kb long. Principal component analysis was performed on the normalized interaction counts for each chromosome, and the scores for the first and second components were recorded in each case.

We used the HiC data to locate the position of centromeres using the method described by Varoquaux (2015). Topologically associated domains (TADs) were estimated from HiC data using TopDom v. 0.0.2 (Shin et al. 2016), setting the minimum number of bins for a domain at 5. We then used Homer v. 4.9 (Heinz et al. 2010) to search for enriched sequence motifs on the edges of TADs, regions designated as ‘gaps’ by TopDom, and in the centromeric windows.

We also tested whether possible mismapping of reads derived from repetitive sequences may have inflated the contact frequencies estimated between repeat-rich sequences. To do this, we produced two distance matrices comparing all AT-rich regions in the genome. In the first matrix, we calculated distances from the number of contacts between regions; in the second, we used the number of repeat families shared by those regions. Finally, we tested for a relationship between repeat content and HiC contacts using the Mantel Test for correlation among distance matrices, as implemented in the R package ade4 v. 1.7-8 (Chessel et al. 2004).

### Influence of genome architecture on gene expression

To examine the degree to which genes within TADs are co-regulated, we used R to fit a linear model with log_2_ fold-change in the expression of a given gene between the in culture and *in planta* RNAseq datasets (obtained from the differential gene expression analysis described above) as the response variable and TAD membership as the only predictor. We compared this model using the Akaike Information Criterion (AIC) to a null model in which only the intercept was fitted.

We tested the degree to which the three-dimensional structure of the genome influenced gene expression by integrating our RNAseq and HiC data. We first identified the 5% of genes with the highest and lowest (non-zero) expression in culture. We calculated the mean number of observed contacts among genomic windows containing each gene set, and determined the significance of the number of interactions by comparing these results with those obtained from 1000 randomly sampled gene sets of the same size. To determine the effect of genomic distance on interactions among gene sets, the same procedure was performed considering only genes at the following non-overlapping genomic distances { [0-5 kb), [5 kb-20 kb), [20 kb-200 kb), [200 kb-1 Mb), >=1 Mb in *cis*, in *trans* }.

Finally, we investigated whether genes likely to be involved in host-interaction were over-represented within and near AT-rich regions. We identified four subsets of *Epichloë* genes that are likely to be involved in host-interactions:

a. Up-regulated genes (the 5% of genes with the highest up-regulation *in planta*);
b. Genes encoding proteins with signal peptides as detected by SignalP v. 4.0 (Petersen et al. 2011);
c. Genes with homology to known polyketide synthase (PKS), non-ribosomal peptide synthase (NRPS), terpene synthases or siderophore-containing proteins or known to catalyse the production of secondary metabolites (Schardl et al. 2013; Fleetwood et al. 2007; Young et al. 2005);
d. Genes with strain-specific expression (with evidence for expression *in planta* in Fl1 but not part of existing transcriptomes for the current reference strain E2368).

The minimum distance between each Fl1 gene and any AT-rich region was found using the bedtools command ‘closest’. We examined whether any of these subsets of genes were over-represented within 5 kb AT-rich regions using logistic regression in R.

### Software and data availability

A software repository providing the scripts uses to produce the computational results described above is available at https://github.com/dwinter/fl1_genome_scripts. Data underlying these analyses have been deposited at the NCBI under Bioproject PRJNA431450.

## Acknowledgements

We thank colleagues for their helpful contributions: Justin O’sullivan (Auckl Matthew Campbell (currently Hokkaido University); Richard Johnson (AgResearch), Tetsuya Chujo and Yonathan Lukito (Massey University). This research was supported by a grant from the Royal Society of New Zealand through a Rutherford Fellowship (RDF-10-MAU-001) to MPC, a Marsden grant (14-MAU-007) to MPC, ARDG and CAY; a Marsden grant (10-MAU-057) to BS; and by the Tertiary Education Commission via a Bio-Protection Research Centre grant to MPC.

## References

Andrey G, Montavon T, Mascrez B, Gonzalez F, Noordermeer D, Leleu M, Trono D, Spitz F, Duboule D. 2013. A switch between topological domains underlies HoxD genes collinearity in mouse limbs. Science. 340:1234167. https://dx.doi.org/10.1126/science.1234167.

Arachevaleta M, Bacon CW, Hoveland CS, Radcliffe DE. 1989. Effect of the tall fescue endophyte on plant response to environmental stress. Agronomy Journal. 81:83. https://dx.doi.org/10.2134/agronj1989.00021962008100010015x.

Bayat F, Mirlohi A, Khodambashi M. 2009. Effects of endophytic fungi on some drought tolerance mechanisms of tall fescue in a hydroponics culture. Russian Journal of Plant Physiology. 56:510–516. https://dx.doi.org/10.1134/S1021443709040104.

Bonev B, Cavalli G. 2016. Organization and function of the 3D genome. Nature Reviews Genetics. 17:661–678. https://dx.doi.org/10.1038/nrg.2016.112.

Braumann I, van den Berg M, Kempken F. 2008. Repeat induced point mutation in two asexual fungi, *Aspergillus niger* and *Penicillium chrysogenum*. Current Genetics. 53:287–297. https://dx.doi.org/10.1007/s00294-008-0185-y.

Burton JN, Liachko I, Dunham MJ, Shendure J. 2014. Species-level deconvolution of metagenome assemblies with Hi-C–based contact probability maps. G3: Genes|Genomes|Genetics. 4:1339–1346. https://dx.doi.org/10.1534/g3.114.011825.

Byrd AD, Schardl CL, Songlin PJ, Mogen KL, Siegel MR. 1990. The β-tubulin gene of *Epichloë typhina* from perennial ryegrass (*Lolium perenne*). Current Genetics. 18:347–354. https://dx.doi.org/10.1007/BF00318216.

Camacho C, Coulouris G, Avagyan V, Ma N, Papadopoulos J, Bealer K, Madden TL. 2009. BLAST+: architecture and applications. BMC Bioinformatics. 10:421. https://dx.doi.org/10.1186/1471-2105-10-421.

Cambareri EB, Jensen BC, Schabtach E, Selker EU. 1989. Repeat-induced G-C to A-T mutations in *Neurospora*. Science (New York, N.Y.). 244:1571–1575.

Chessel D, Dufour AB, Thioulouse J. 2004. The ade4 package - One-table methods. R News. 4:5–10.

Christensen MJ. 1996. Antifungal activity in grasses infected with *Acremonium* and *Epichloë* endophytes. Australasian Plant Pathology. 25:186–191. https://dx.doi.org/10.1071/AP96032.

Chuong EB, Elde NC, Feschotte C. 2017. Regulatory activities of transposable elements: from conflicts to benefits. Nature Reviews Genetics. 18:71–86. https://dx.doi.org/10.1038/nrg.2016.139.

Clarke BB, Belanger FC, Ambrose KV, Wang R, Tian Z. 2017. The *Epichloë festucae* antifungal protein has activity against the plant pathogen *Sclerotinia homoeocarpa*, the causal agent of dollar spot disease. Scientific Reports. 7:5643. https://dx.doi.org/10.1038/s41598-017-06068-4.

Clay K, Cheplick GP. 1989. Effect of ergot alkaloids from fungal endophyte-infected grasses on fall armyworm (*Spodoptera frugiperda*). Journal of Chemical Ecology. 15:169–182. https://dx.doi.org/10.1007/BF02027781.

Crénès G, Moundras C, Demattei M-V, Bigot Y, Petit A, Renault S. 2010. Target site selection by the mariner-like element, Mos1. Genetica. 138:509–517. https://dx.doi.org/10.1007/s10709-009-9387-6.

Cubeñas-Potts C, Rowley MJ, Lyu X, Li G, Lei EP, Corces VG. 2017. Different enhancer classes in Drosophila bind distinct architectural proteins and mediate unique chromatin interactions and 3D architecture. Nucleic Acids Research. 45:1714–1730. https://dx.doi.org/10.1093/nar/gkw1114.

Dixon JR, Selvaraj S, Yue F, Kim A, Li Y, Shen Y, Hu M, Liu JS, Ren B. 2012. Topological domains in mammalian genomes identified by analysis of chromatin interactions. Nature. 485:376–380. https://dx.doi.org/10.1038/nature11082.

Dobin A, Davis CA, Schlesinger F, Drenkow J, Zaleski C, Jha S, Batut P, Chaisson M, Gingeras TR. 2013. STAR: ultrafast universal RNA-seq aligner. Bioinformatics. 29:15–21. https://dx.doi.org/10.1093/bioinformatics/bts635.

Dong S, Raffaele S, Kamoun S. 2015. The two-speed genomes of filamentous pathogens: waltz with plants. Current Opinion in Genetics & Development. 35:57–65. https://dx.doi.org/10.1016/j.gde.2015.09.001.

Duan Z, Andronescu M, Schutz K, Mcllwain S, Kim YJ, Lee C, Shendure J, Fields S, Blau CA, Noble WS. 2010. A Three-Dimensional Model of the Yeast Genome. Nature. 465:363–367. https://dx.doi.org/10.1038/nature08973.

Eaton CJ, Cox MP, Ambrose B, Becker M, Hesse U, Schardl CL, Scott B. 2010. Disruption of signaling in a fungal-grass symbiosis leads to pathogenesis. Plant Physiology. 153:1780–1794. https://dx.doi.org/10.1104/pp.110.158451.

Eaton CJ, Cox MP, Scott B. 2011. What triggers grass endophytes to switch from mutualism to pathogenism? Plant Science. 180:190–195. https://dx.doi.org/10.1016/j.plantsci.2010.10.002.

Eaton CJ, Dupont P-Y, Solomon P, Clayton W, Scott B, Cox MP. 2015. A core gene set describes the molecular basis of mutualism and antagonism in *Epichloë* spp. Molecular plant-microbe interactions: MPMI. 28:218–231. https://dx.doi.org/10.1094/MPMI-09-14-0293-FI.

Edelman LB, Fraser P. 2012. Transcription factories: genetic programming in three dimensions. Current Opinion in Genetics & Development. 22:110–114. https://dx.doi.org/10.1016/j.gde.2012.01.010.

Edgar RC. 2004. MUSCLE: multiple sequence alignment with high accuracy and high throughput. Nucleic Acids Research. 32:1792–1797. https://dx.doi.org/10.1093/nar/gkh340.

Eser U, Chandler-Brown D, Ay F, Straight AF, Duan Z, Noble WS, Skotheim JM. 2017. Form and function of topologically associating genomic domains in budding yeast. Proceedings of the National Academy of Sciences. 114:E3061–E3070. https://dx.doi.org/10.1073/pnas.1612256114.

Faino L, Seidl MF, Shi-Kunne X, Pauper M, Berg GCM van den, Wittenberg AHJ, Thomma BPHJ. 2016. Transposons passively and actively contribute to evolution of the two-speed genome of a fungal pathogen. Genome Research. gr.204974.116. https://dx.doi.org/10.1101/gr.204974.116.

Feschotte C, Zhang X, Wessler SR. 2002. Miniature inverted-repeat transposable elements and their relationship to established DNA transposons. In: Mobile DNA II. pp. 1147–1158. http://www.asmscience.org/content/book/10.1128/9781555817954.chap50 (Accessed November 23, 2017).

Fleetwood DJ, Khan AK, Johnson RD, Young CA, Mittal S, Wrenn RE, Hesse U, Foster SJ, Schardl CL, Scott B. 2011. Abundant degenerate miniature inverted-repeat transposable elements in genomes of epichloid fungal endophytes of grasses. Genome Biology and Evolution. 3:1253–1264. https://dx.doi.org/10.1093/gbe/evr098.

Fleetwood DJ, Scott B, Lane GA, Tanaka A, Johnson RD. 2007. A complex ergovaline gene cluster in *Epichloë* endophytes of grasses. Applied and Environmental Microbiology. 73:2571–2579. https://dx.doi.org/10.1128/AEM.00257-07.

Funabiki H, Hagan I, Uzawa S, Yanagida M. 1993. Cell cycle-dependent specific positioning and clustering of centromeres and telomeres in fission yeast. The Journal of Cell Biology. 121:961–976.

Gladyshev E. 2017. Repeat-induced point mutation (RIP) and other genome defense mechanisms in fungi. Microbiology spectrum. 5. https://dx.doi.org/10.1128/microbiolspec.FUNK-0042-2017.

Gladyshev E, Kleckner N. 2017. DNA sequence homology induces cytosine-to-thymine mutation by a heterochromatin-related pathway in *Neurospora*. Nature Genetics. 49:ng.3857. https://dx.doi.org/10.1038/ng.3857.

Gonzalez-Sandoval A, Gasser SM. 2016. On TADs and LADs: spatial control over gene expression. Trends in Genetics. 32:485–495. https://dx.doi.org/10.1016/j.tig.2016.05.004.

Grand RS, Pichugina T, Gehlen LR, Jones MB, Tsai P, Allison JR, Martienssen R, O’Sullivan JM. 2014. Chromosome conformation maps in fission yeast reveal cell cycle dependent sub nuclear structure. Nucleic Acids Research. 42:12585–12599. https://dx.doi.org/10.1093/nar/gku965.

Heinz S, Benner C, Spann N, Bertolino E, Lin YC, Laslo P, Cheng JX, Murre C, Singh H, Glass CK. 2010. Simple combinations of lineage-determining transcription factors prime cis-regulatory elements required for macrophage and B cell identities. Molecular Cell. 38:576–589. https://dx.doi.org/10.1016/j.molcel.2010.05.004.

Hou C, Zhao H, Tanimoto K, Dean A. 2008. CTCF-dependent enhancer-blocking by alternative chromatin loop formation. Proceedings of the National Academy of Sciences of the United States of America. 105:20398–20403. https://dx.doi.org/10.1073/pnas.0808506106.

Kauppinen M, Saikkonen K, Helander M, Pirttilä AM, Wäli PR. 2016. *Epichloë* grass endophytes in sustainable agriculture. Nature Plants. 2:15224. https://dx.doi.org/10.1038/nplants.2015.224.

Kimmons CA. 1990. Nematode reproduction on endophyte-infected and endophyte-free tall fescue. Plant Disease. 74:757. https://dx.doi.org/10.1094/PD-74-0757.

Klocko AD, Ormsby T, Galazka JM, Leggett NA, Uesaka M, Honda S, Freitag M, Selker EU. 2016. Normal chromosome conformation depends on subtelomeric facultative heterochromatin in *Neurospora* crassa. Proceedings of the National Academy of Sciences. 113:15048–15053. https://dx.doi.org/10.1073/pnas.1615546113.

Koren S, Walenz BP, Berlin K, Miller JR, Bergman NH, Phillippy AM. 2017. Canu: scalable and accurate long-read assembly via adaptive k-mer weighting and repeat separation. bioRxiv. 071282. https://dx.doi.org/10.1101/071282.

Leuchtmann A. 1994. Isozyme relationships of *Acremonium* endophytes from twelve *Festuca* species. Mycological Research. 98:25–33. https://dx.doi.org/10.1016/S0953-7562(09)80331-6.

Li G, Ruan X, Auerbach RK, et al. 2012. Extensive promoter-centered chromatin interactions provide a topological basis for transcription regulation. Cell. 148:84–98. https://dx.doi.org/10.1016/j.cell.2011.12.014.

Li H. 2013. Aligning sequence reads, clone sequences and assembly contigs with BWA-MEM. https://arxiv.org/abs/1303.3997# (Accessed February 28, 2017).

Lieberman-Aiden E, van Berkum NL, Williams L, Imakaev M, Ragoczy T, Telling A, Amit I, Lajoie BR, Sabo PJ, Dorschner MO, Sandstrom R, Bernstein B, Bender MA, Groudine M, Gnirke A, Stamatoyannopoulos J, Mirny LA, Lander ES, Dekker J. 2009. Comprehensive mapping of long-range interactions reveals folding principles of the human genome. Science (New York, N.Y.). 326:289–293. https://dx.doi.org/10.1126/science.1181369.

Love MI, Huber W, Anders S. 2014. Moderated estimation of fold change and dispersion for RNA-seq data with DESeq2. Genome Biology. 15:550. https://dx.doi.org/10.1186/s13059-014-0550-8.

Lugtenberg BJJ, Caradus JR, Johnson LJ. 2016. Fungal endophytes for sustainable crop production. FEMS microbiology ecology. 92. https://dx.doi.org/10.1093/femsec/fiw194.

Malinowski DP, Belesky DP. 2000. Adaptations of endophyte-infected cool-season grasses to environmental stresses: mechanisms of drought and mineral stress tolerance. Crop Science. 40:923–940. https://dx.doi.org/10.2135/cropsci2000.404923x.

Margolin BS, Garrett-Engele PW, Stevens JN, Fritz DY, Garrett-Engele C, Metzenberg RL, Selker EU. 1998. A methylated Neurospora 5S rRNA pseudogene contains a transposable element inactivated by repeat-induced point mutation. Genetics. 149:1787–1797.

Mizuguchi T, Fudenberg G, Mehta S, Belton J-M, Taneja N, Folco HD, FitzGerald P, Dekker J, Mirny L, Barrowman J, Grewal SIS. 2014. Cohesin-dependent globules and heterochromatin shape 3D genome architecture in *S. pombe*. Nature. 516:432–435. https://dx.doi.org/10.1038/nature13833.

Mondo SJ, Dannebaum RO, Kuo RC, et al. 2017. Widespread adenine N6-methylation of active genes in fungi. Nature Genetics. 49:964–968. https://dx.doi.org/10.1038/ng.3859.

Moon CD, Tapper BA, Scott B. 1999. Identification of *Epichloë* endophytes in planta by a microsatellite-based pcr fingerprinting assay with automated analysis. Applied and Environmental Microbiology. 65:1268–1279.

Nicodemi M, Pombo A. 2014. Models of chromosome structure. Current Opinion in Cell Biology. 28:90–95. https://dx.doi.org/10.1016/j.ceb.2014.04.004.

Nora EP, Lajoie BR, Schulz EG, Giorgetti L, Okamoto I, Servant N, Piolot T, van Berkum NL, Meisig J, Sedat J, Gribnau J, Barillot E, Blüthgen N, Dekker J, Heard E. 2012. Spatial partitioning of the regulatory landscape of the X-inactivation centre. Nature. 485:381–385. https://dx.doi.org/10.1038/nature11049.

Paradis E, Claude J, Strimmer K. 2004. APE: analyses of phylogenetics and evolution in R language. Bioinformatics. 20:289–290. https://dx.doi.org/10.1093/bioinformatics/btg412.

Petersen TN, Brunak S, von Heijne G, Nielsen H. 2011. SignalP 4.0: discriminating signal peptides from transmembrane regions. Nature Methods. 8:785–786. https://dx.doi.org/10.1038/nmeth.1701.

Quinlan AR, Hall IM. 2010. BEDTools: a flexible suite of utilities for comparing genomic features. Bioinformatics. 26:841–842. https://dx.doi.org/10.1093/bioinformatics/btq033.

R Core Team 2017. R: a language and environment for statistical computing. Vienna, Austria http://www.R-project.org.

Roberts A, Pimentel H, Trapnell C, Pachter L. 2011. Identification of novel transcripts in annotated genomes using RNA-Seq. Bioinformatics. 27:2325–2329. https://dx.doi.org/10.1093/bioinformatics/btr355.

Rowan DD, Gaynor DL. 1986. Isolation of feeding deterrents against argentine stem weevil from ryegrass infected with the endophyte *Acremonium loliae*. Journal of Chemical Ecology. 12:647–658. https://dx.doi.org/10.1007/BF01012099.

Saikkonen K, Young CA, Helander M, Schardl CL. 2016. Endophytic *Epichloë* species and their grass hosts: from evolution to applications. Plant Molecular Biology. 90:665–675. https://dx.doi.org/10.1007/s11103-015-0399-6.

Sanyal K, Baum M, Carbon J. 2004. Centromeric DNA sequences in the pathogenic yeast *Candida albicans* are all different and unique. Proceedings of the National Academy of Sciences of the United States of America. 101:11374–11379. https://dx.doi.org/10.1073/pnas.0404318101.

Schardl CL. 2001. *Epichloë festucae* and related mutualistic symbionts of grasses. Fungal Genetics and Biology. 33:69–82. https://dx.doi.org/10.1006/fgbi.2001.1275.

Schardl CL, Grossman RB, Nagabhyru P, Faulkner JR, Mallik UP. 2007. Loline alkaloids: Currencies of mutualism. Phytochemistry. 68:980–996. https://dx.doi.org/10.1016/j.phytochem.2007.01.010.

Schardl CL, Young CA, Hesse U, et al. 2013. Plant-symbiotic fungi as chemical engineers: multi-genome analysis of the Cavicipitaceae reveals dynamics of alkaloid loci. PLOS Genetics. 9:e1003323. https://dx.doi.org/10.1371/journal.pgen.1003323.

Schatz MC, Delcher AL, Salzberg SL. 2010. Assembly of large genomes using second-generation sequencing. Genome Research. 20:1165–1173. https://dx.doi.org/10.1101/gr.101360.109.

Schmid MW, Grob S, Grossniklaus U. 2015. HiCdat: a fast and easy-to-use Hi-C data analysis tool. BMC Bioinformatics. 16:277. https://dx.doi.org/10.1186/s12859-015-0678-x.

Schmidt SM, Houterman PM, Schreiver I, Ma L, Amyotte S, Chellappan B, Boeren S, Takken FLW, Rep M. 2013. MITEs in the promoters of effector genes allow prediction of novel virulence genes in *Fusarium oxysporum*. BMC Genomics. 14:119. https://dx.doi.org/10.1186/1471-2164-14-119.

Scott B, Becker Y, Becker M, Cartwright G. 2012. Morphogenesis, Growth, and Development of the Grass Symbiont *Epichlöe festucae*. In: Morphogenesis and Pathogenicity in Fungi.Topics in Current Genetics Springer, Berlin, Heidelberg pp. 243–264. https://dx.doi.org/10.1007/978-3-642-22916-9_12.

Scott B, Wrenn RE, May KJ, Takemoto D, Young CA, Tanaka A, Fleetwood DJ, Johnson RD. 2010. Regulation and Functional Analysis of Bioprotective Metabolite Genes from the Grass Symbiont Epichloe festucae. In: Recent developments in management of plant diseases. Gisi, U, Chet, I, & Gullino, ML, editors. Springer Netherlands: Dordrecht pp. 199–213. https://dx.doi.org/10.1007/978-1-4020-8804-9_15.

Shin H, Shi Y, Dai C, Tjong H, Gong K, Alber F, Zhou XJ. 2016. TopDom: an efficient and deterministic method for identifying topological domains in genomes. Nucleic Acids Research. 44:e70–e70. https://dx.doi.org/10.1093/nar/gkv1505.

Smit AF, Hubley R, Green P. 1996. RepeatMasker Open-3.0.

Stamatakis A. 2006. RAxML-VI-HPC: maximum likelihood-based phylogenetic analyses with thousands of taxa and mixed models. Bioinformatics. 22:2688.

Tanaka A, Christensen MJ, Takemoto D, Park P, Scott B. 2006. Reactive oxygen species play a role in regulating a fungus–perennial ryegrass mutualistic interaction. The Plant Cell. 18:1052–1066. https://dx.doi.org/10.1105/tpc.105.039263.

Tanaka A, Takemoto D, Chujo T, Scott B. 2012. Fungal endophytes of grasses. Current Opinion in Plant Biology. 15:462–468. https://dx.doi.org/10.1016/j.pbi.2012.03.007.

Testa AC, Oliver RP, Hane JK. 2016. OcculterCut: a comprehensive survey of AT-rich regions in fungal genomes. Genome Biology and Evolution. 8:2044–2064. https://dx.doi.org/10.1093/gbe/evw121.

Trapnell C, Roberts A, Goff L, Pertea G, Kim D, Kelley DR, Pimentel H, Salzberg SL, Rinn JL, Pachter L. 2012. Differential gene and transcript expression analysis of RNA-seq experiments with TopHat and Cufflinks. Nature Protocols. 7:562–578. https://dx.doi.org/10.1038/nprot.2012.016.

Ulianov SV, Khrameeva EE, Gavrilov AA, Flyamer IM, Kos P, Mikhaleva EA, Penin AA, Logacheva MD, Imakaev MV, Chertovich A, Gelfand MS, Shevelyov YY, Razin SV. 2015. Active chromatin and transcription play a key role in chromosome partitioning into topologically associating domains. Genome Research. gr.196006.115. https://dx.doi.org/10.1101/gr.196006.115.

Varoquaux N, Liachko I, Ay F, Burton JN, Shendure J, Dunham MJ, Vert J-P, Noble WS. 2015. Accurate identification of centromere locations in yeast genomes using Hi-C. Nucleic Acids Research. 43:5331–5339. https://dx.doi.org/10.1093/nar/gkv424.

Walker BJ, Abeel T, Shea T, Priest M, Abouelliel A, Sakthikumar S, Cuomo CA, Zeng Q, Wortman J, Young SK, Earl AM. 2014. Pilon: an integrated tool for comprehensive microbial variant detection and genome assembly improvement. PLOS ONE. 9:e112963. https://dx.doi.org/10.1371/journal.pone.0112963.

Wei L, Xiao M, An Z, Ma B, Mason AS, Qian W, Li J, Fu D. 2013. New insights into nested long terminal repeat retrotransposons in *Brassica* species. Molecular Plant. 6:470–482. https://dx.doi.org/10.1093/mp/sss081.

Young CA, Bryant MK, Christensen MJ, Tapper BA, Bryan GT, Scott B. 2005. Molecular cloning and genetic analysis of a symbiosis-expressed gene cluster for lolitrem biosynthesis from a mutualistic endophyte of perennial ryegrass. Molecular Genetics and Genomics. 274:13–29. https://dx.doi.org/10.1007/s00438-005-1130-0.

Zhang Y, McCord RP, Ho Y-J, Lajoie BR, Hildebrand DG, Simon AC, Becker MS, Alt FW, Dekker J. 2012. Chromosomal translocations are guided by the spatial organization of the genome. Cell. 148:908–921. https://dx.doi.org/10.1016/j.cell.2012.02.002.

